# Core NHEJ Factors Maintain Synapsis Across Diverse DNA End Structures

**DOI:** 10.64898/2026.09.30.755744

**Authors:** Brandon C. Case, Isabel T. Burger, Joseph J. Loparo

## Abstract

DNA double strand breaks (DSBs) generated by genotoxic stress frequently contain chemically diverse termini that require end processing before ligation by non-homologous end joining (NHEJ). How end chemistry influences assembly and persistence of NHEJ synaptic complexes remains unclear. Using single-molecule imaging approaches in *Xenopus* egg extract, we measured formation and stability of the long-range (LRC) and short-range (SRC) complexes on nine DNA substrates spanning kinase-, polymerase- and nuclease-dependent processing requirements. LRC formation and lifetime were comparable across end types, and SRC formation and stability were largely insensitive to end chemistry. DNA hairpins, physiological intermediates of V(D)J recombination, were a notable exception that formed LRCs but inefficiently transitioned to SRCs. Depletion of cognate processing factors had modest effects on SRC formation and persistence, indicating that core NHEJ factors provide the principal structural support for synapsis across diverse termini. Although XLF was required for SRC formation on all substrates, reducing an XLF dimer from two intact tail/Ku-binding motifs to one selectively reduced productive SRC formation on ends requiring processing. Together, these results support a model in which partially redundant protein-protein interactions between core NHEJ factors maintain synapsis while end-specific processing reactions shape repair outcome.

## Introduction

The failure to repair DNA double strand breaks (DSBs) can result in genomic instability, tumorigenesis and cell death. DSBs are repaired primarily by two pathways: homologous recombination (HR) and non-homologous end joining (NHEJ). HR operates predominantly in S and G2 phase, when a sister chromatid is available as a repair template, while NHEJ functions throughout the cell cycle and repairs the majority of DSBs in mammalian cells by directly rejoining the two broken DNA ends.^1–4^

Although NHEJ is often described as a direct ligation of DNA ends, DSBs generated by ionizing radiation, oxidative stress and genotoxic chemotherapeutics frequently contain termini that cannot be ligated immediately.^5–8^ These breaks can contain abnormal terminal phosphates, damaged nucleotides, short gaps or overhangs, flaps and other end structures that require processing before ligation.^9^ In addition, programmed DSBs formed during V(D)J recombination generate hairpin termini that must be opened prior to joining.^10^ A diverse group of processing enzymes act on these termini including tyrosyl-DNA phosphodiesterase 1 (TDP1), polynucleotide kinase phosphatase (PNKP), DNA polymerases λ and μ, and Artemis.^9^ Because end processing alters the repair junction, NHEJ must coordinate processing with ligation to limit unnecessary or excessive modification of broken ends.^11–17^

A central feature of NHEJ is the formation of a synaptic complex that tethers the DNA ends together and prevents their mispairing with other breaks.^4^ Synapsis proceeds through a long-range and a short-range complex (LRC and SRC), which differ in the distance between the DNA ends.^18,19^ The Ku70/Ku80 heterodimer (Ku) rapidly recognizes DNA ends and recruits DNA-dependent protein kinase catalytic subunit (DNA-PKcs) to form the DNA-PK holoenzyme.^20–22^ Interactions between DNA-PK complexes on opposing ends tether the break in the LRC, with PAXX providing an additional bridge through interactions with Ku.^18,19,23^ Subsequent DNA-PKcs phosphorylation as well as recruitment and rearrangement of core NHEJ factors, including XRCC4, DNA ligase IV (LigIV) and XLF, bring the ends into close alignment in the SRC.^24–29^ Multiple protein-mediated contacts span the break in this complex, and processing is thought to occur within the SRC between ligation attempts.^4,16,17,27,29^

Much of our current mechanistic understanding of NHEJ synapsis derives from studies of readily ligatable DNA ends.^4^ Single-molecule studies on non-ligatable termini have concluded that end structure may promote interaction with LigIV or SRC formation.^30,31^ However, it has remained difficult to determine how end chemistry influences synapsis in systems that retain the DNA-PKcs-dependent LRC-to-SRC transition and the endogenous end processing activities to repair these diverse termini. DNA end structure could alter the formation or stability of the LRC and SRC, affect progression between these states or recruit processing enzymes that themselves may stabilize or destabilize the complex. Alternatively, the core NHEJ machinery may form a scaffold that is broadly tolerant of diverse termini, allowing processing enzymes to act within an already assembled synaptic complex. In this model, the contribution of individual bridging interactions may become more important when ligation is delayed, and the DNA ends must remain synapsed during processing.

To distinguish between these possibilities, we used single-molecule imaging in *Xenopus* egg extract to measure LRC and SRC formation and stability on nine DNA end structures with distinct processing requirements. *Xenopus* egg extract contains a broad complement of endogenous proteins, including core NHEJ factors and DNA end-processing enzymes defined in human cells.^12,32–34^ It robustly joins both ligation-competent and processing-dependent DNA ends, making it well suited for mechanistic analysis of NHEJ synapsis.^16,18^ We find that synapsis is largely unaffected by end chemistry across non-hairpin substrates and is maintained independently of cognate processing factors. Hairpin ends constitute a notable exception, forming LRCs but inefficiently transitioning to SRCs. We further tested the contribution of XLF, a central component of the SRC bridging network. Although XLF is broadly required for SRC formation, reducing an XLF dimer from two intact tail/Ku-binding motifs to one selectively impaired productive SRC formation on substrates requiring end processing. Together, these findings support a model in which a partially redundant network of protein-protein interactions maintains synapsis across varied DNA end structures, while specific bridging interactions become especially important when processing delays ligation. This model provides a framework for how NHEJ maintains end proximity while repairing the chemically diverse termini generated by genotoxic and physiological DSBs.

## Results

### Long-range complex formation and stability

To determine how DNA end structure impacts LRC formation and stability, we used a previously developed single-molecule TIRF assay in which DNA in solution colocalizes with surface-tethered DNA substrates (Figure 1A).^18^ In *Xenopus laevis* high-speed supernatant (HSS) egg extract, colocalization of the two DNA molecules is dependent on Ku and DNA-PKcs, highlighting the assay’s specificity to NHEJ.^18^ In the assay, green laser excitation images a stable fluorescence signal as the DNA is stably bound to the surface (Figure 1B, top) while red laser excitation shows colocalization of the in-solution DNAs when they are transiently synapsed with surface DNAs (Figure 1B, bottom). Individual molecules are represented in a rastergram as a line over time with colocalization events sorted based on the time to first binding and shaded according to their dwell time (Supplementary Figure 1A). Transition from the LRC to the SRC is marked by the appearance of FRET as the DNA ends move into close alignment, though transition to the SRC is inefficient in the two-piece assay (data not shown).^18,35^ To understand how different DNA ends affect LRC formation and stability, oligo pairs forming different structures were ligated onto backbones to create a variety of DNA end structures (Figure 1C, Supplementary Table 1).

**Figure 1:**
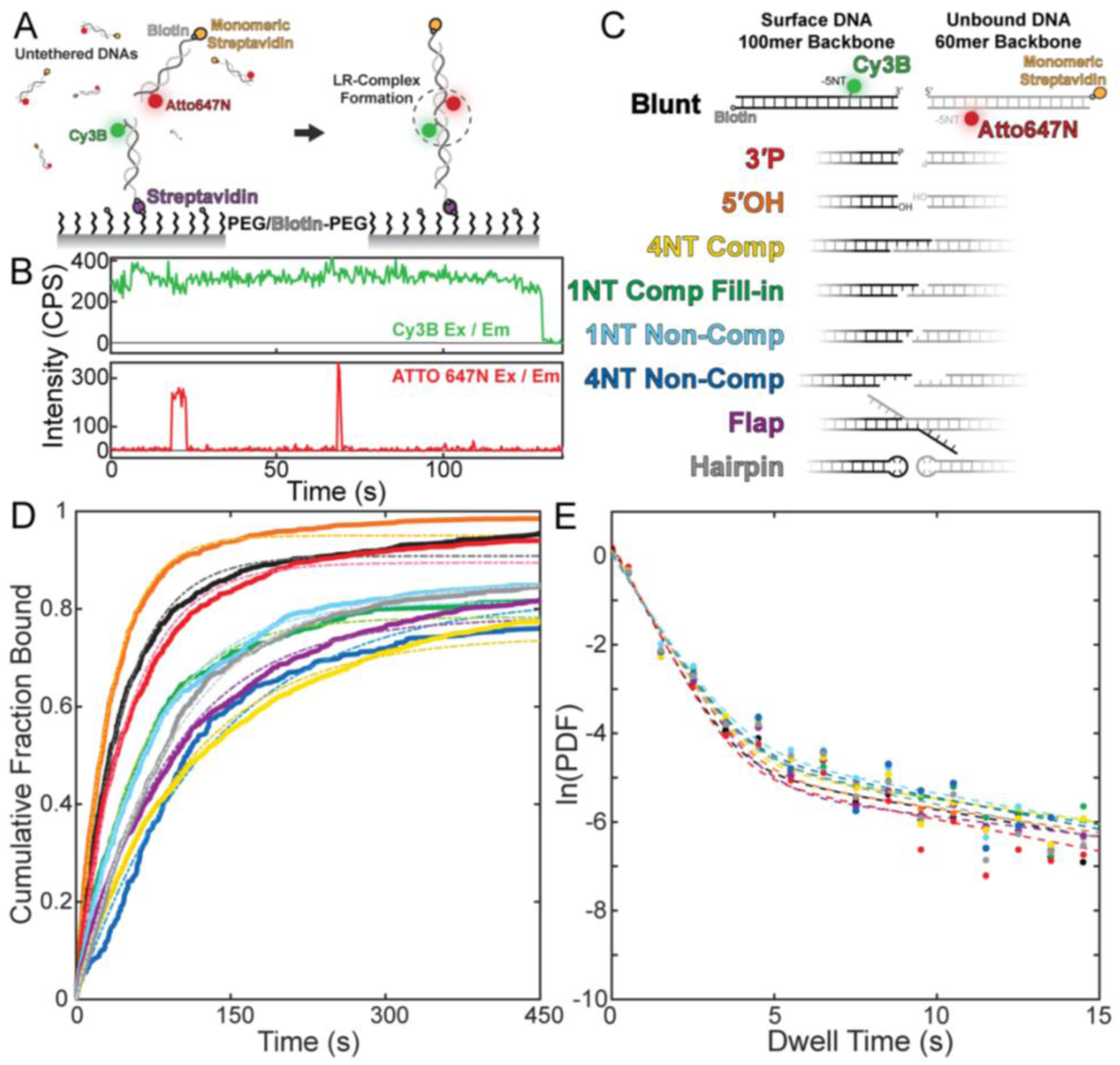
DNA end structure does not influence LRC formation or stability. (A) Schematic of the two-piece single-molecule colocalization assay to monitor LRC formation and stability with DNA colocalization shown as a dashed circle. (B) Representative single-molecule trace showing stable Cy3B fluorescence from a surface DNA (upper) and transient colocalization of an in-solution ATTO 647N DNA (lower). (C) Schematic of the DNA substrate backbones and different DNA end structures used in the two-piece assay. (D) Cumulative fraction of colocalized molecules over time for different DNA end types, rastergrams shown in Supplementary Figure 1A; time to first binding events were fit to a single exponential model, fits shown in Table 1. (E) Dwell time histograms of individual DNA binding events were best described by a double exponential decay and plotted on a natural log scale to compare the relative stability on each DNA substrate; histograms shown in Supplementary Figure 1B, fits in Table 2.

**Table 1.** Two-piece DNA LRC Binding Rates.

| DNA | Amp (%)<br>[95% CI] | $k_{obs}$ (s<sup>-1</sup>)<br>[95% CI] | N |
| --- | --- | --- | --- |
| Blunt | 90.9 [90.5 91.2] | 0.0237 [0.0235 0.0240] | 791 |
| 3'P | 89.5 [89.1 89.9] | 0.0208 [0.0205 0.0211] | 819 |
| 5'OH | 95.1 [94.7 95.4] | 0.0312 [0.0308 0.0315] | 702 |
| 4NT Comp | 74.4 [74.0 74.8] | 0.0098 [0.0096 0.0099] | 1953 |
| 1NT Comp Fill-in | 78.4 [77.7 79.0] | 0.0165 [0.0162 0.0168] | 595 |
| 1NT Non-Comp | 81.4 [80.8 82.1] | 0.0146 [0.0144 0.0149] | 578 |
| 4NT Non-Comp | 82.6 [81.5 83.6] | 0.0076 [0.0075 0.0078] | 497 |
| Flap | 78.3 [77.9 78.7] | 0.0112 [0.0110 0.0113] | 1495 |
| Hairpin | 85.5 [84.9 86.2] | 0.0105 [0.0103 0.0107] | 640 |

**Table 2.** Two-piece DNA LRC Colocalization Dwell Times.

| DNA End | Amp<sub>1</sub> (%)<br>[95% CI] | $\tau_1$ (%)<br>[95% CI] | Amp<sub>2</sub> (%)<br>[95% CI] | $\tau_2$ (%)<br>[95% CI] | N |
| --- | --- | --- | --- | --- | --- |
| Blunt | 91 [90 92] | 0.74 [0.72 0.76] | 9 [8 10] | 7.9 [6.9 9.1] | 7985 |
| 3'P | 92 [91 94] | 0.71 [0.68 0.73] | 8 [6 9] | 7.1 [5.6 9.5] | 6807 |
| 5'OH | 91 [89 92] | 0.80 [0.77 0.83] | 9 [8 11] | 9.1 [7.4 11.3] | 9330 |
| 4NT Comp | 89 [87 90] | 0.80 [0.76 0.83] | 11 [10 13] | 9.6 [7.8 11.6] | 6778 |
| 1NT Comp Fill-in | 89 [86 91] | 0.86 [0.81 0.91] | 11 [9 14] | 8.7 [6.8 10.9] | 2514 |
| 1NT Non-Comp | 87 [84 89] | 0.87 [0.82 0.92] | 13 [11 16] | 7.6 [6.2 9.3] | 2870 |
| 4NT Non-Comp | 87 [83 90] | 0.81 [0.76 0.88] | 13 [10 17] | 7.0 [5.0 9.3] | 2153 |
| Flap | 92 [90 94] | 0.82 [0.78 0.86] | 8 [6 10] | 11.3 [7.5 16.4] | 9176 |
| Hairpin | 89 [86 92] | 0.80 [0.76 0.85] | 11 [8 14] | 6.6 [4.9 8.9] | 4692 |

We constructed nine different DNA substrates containing ends with distinct end processing requirements that were previously shown to undergo the expected processing reactions in egg extract by sequencing of repair junctions.^16^ These included blunt ends and four nucleotide complementary overhangs (4NT Comp) that can be directly ligated, 3ʹP and 5ʹOH ends, which require processing by PNKP, two nucleotide ssDNA overhangs that contain one nucleotide of terminal complementarity (1NT Comp Fill-in) that are preferentially filled in by Pol λ, one and four nucleotide ssDNA overhangs that are non-complementary (1NT Non-Comp and 4NT Non-Comp, respectively) and are preferentially processed by Pol μ, as well as flap and hairpin containing DNAs that are processed by nucleases (Figure 1C; oligos listed in Supplementary Table 1). To measure the rate of LRC formation, we immobilized individual DNA substrates on the surface and flowed in the corresponding DNA in HSS extract.

Binding of in-solution DNAs to surface DNAs resulted in an increase in the cumulative fraction of colocalized molecules bound over time (Figure 1D). Unbound blunt DNA ends, shown in black, underwent rapid and robust colocalization with an observed binding rate, *k_obs_*, of 0.024 s^−1^ with 91% of surface DNAs registering at least one binding event in the 7.5 min observation window (Figure 1D, Table 1). All nine DNA substrates showed comparable rates of LRC formation with binding rates ranging from 0.008 to 0.03 s^−1^, and surface DNAs showing colocalization to more than 74% of molecules (Figure 1D, Table 1).

Next, we asked whether DNA end structures influence the stability of the LRC. Using the same single-molecule assay, we compared the stability of complexes by fitting the distribution of individual colocalization lifetimes to obtain a dwell time. For blunt DNA ends, the dwell time distribution was best fit to a double-exponential model in which the first population accounts for 91% of events with a dwell time of 0.74 s and the second population accounts for 9% of events with a dwell time of 7.9 s, which is comparable to previously reported data (Figure 1E, Supplementary Figure 1B, Table 2). These two phases likely correspond to LRCs with heterogeneous protein compositions, perhaps lacking PAXX in the more transient population.^23^ As observed with the LRC formation rate, all nine DNA end structures exhibited comparable dwell times. The short-lived population accounted for 87 – 92% of events with dwell times ranging from 0.7 – 0.9 s and the stable population accounted for 8 – 13% of events with dwell times ranging from 6.6 – 11.2 s (Figure 1E, Supplementary Figure 1B, Table 2). Overall, these data show that LRC formation and stability are independent of DNA end structure, which is consistent with the DNA ends being held apart from each other by DNA-PK dimerization.

### Short-range complex formation

We next asked how DNA end structure impacts formation of the SRC in which the DNA ends are brought into close alignment.^18^ To increase the efficiency of SRC formation, we used a previously established single-molecule FRET DNA circularization assay in which the donor and acceptor fluorophores are present on opposing ends of the same 3 kb surface-tethered DNA molecule (Figure 2A, C).^16–18^ Because the circularization substrate maintains the two ends on the same molecule, resulting in a high local concentration, SRC formation is substantially more frequent than in the two-piece assay.^18^ Using the same DNA ends from the single-molecule LRC assay, we asked how SRC formation, observed as an increase in FRET, is affected by various DNA structures.

**Figure 2:**
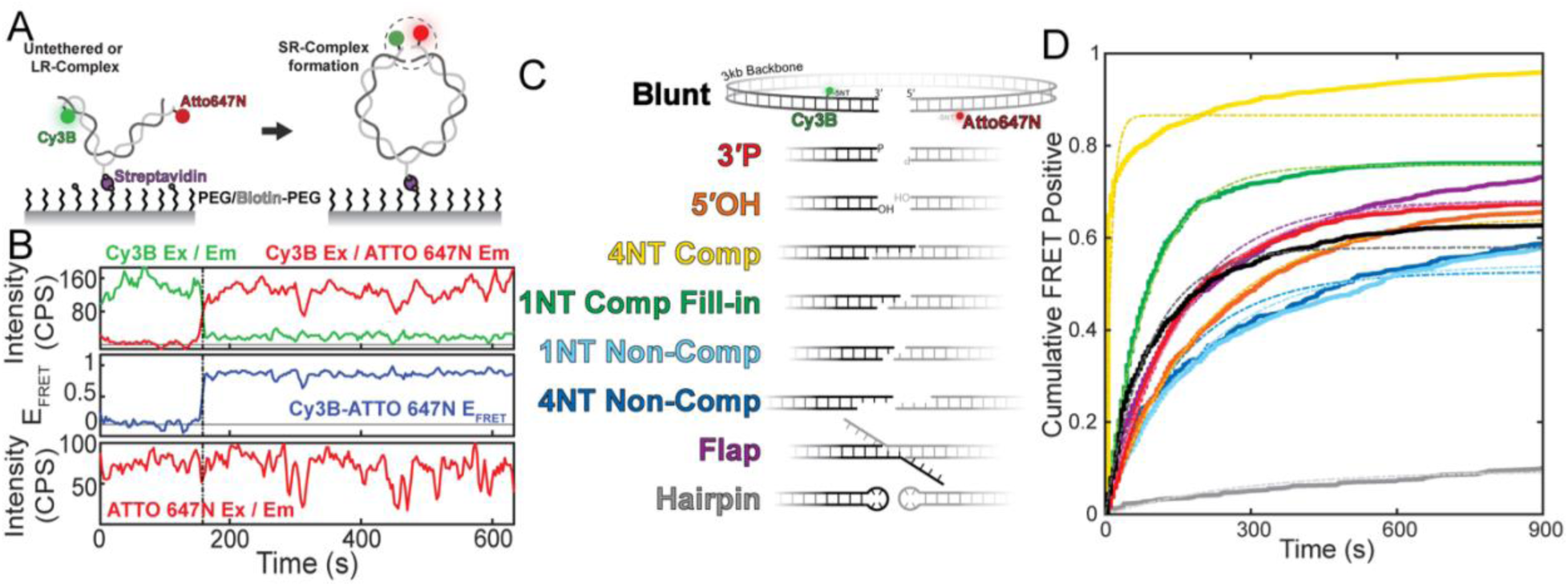
DNA end structure does not influence SRC formation except for hairpin ends. (A) Schematic of the smFRET circularization assay to monitor SRC formation and stability with FRET formation shown as a dashed circle. (B) Representative single-molecule trace shows stable DNA on the surface from Cy3B (upper) or ATTO 647N (lower) excitation and emission. At the grey dashed line, a concomitant decrease in Cy3B emission and increase in ATTO 647N emission upon Cy3B excitation (upper) are indicative of FRET (middle). (C) Schematic of the different DNA substrates used in the smFRET assay. (D) Cumulative fraction of FRET positive molecules, sorted by time to first FRET, shows an exponential increase over time for the different DNA substrates; rastergrams shown in Supplementary Figure 2B. Time to first binding events were fit to a single exponential model, fits shown in Table 3.

**Table 3.** DNA Circularization SRC Formation Rates for ^WT^HSS Extract.

| DNA | Amp (%)<br>[95% CI] | $k_{obs}$ (s<sup>-1</sup>)<br>[95% CI] | N |
| --- | --- | --- | --- |
| Blunt | 57.9 [57.4 58.4] | 0.0109 [0.0107 0.0111] | 1782 |
| 3'P | 68.0 [67.4 68.7] | 0.0062 [0.0061 0.0063] | 1415 |
| 5'OH | 64.6 [64.0 65.2] | 0.0049 [0.0048 0.0050] | 1731 |
| 4NT Comp | 86.7 [86.5 86.9] | 0.0868 [0.0862 0.0873] | 2030 |
| 1NT Comp Fill-in | 75.8 [75.2 76.5] | 0.0112 [0.0110 0.0114] | 1157 |
| 1NT Non-Comp | 54.2 [53.7 54.6] | 0.0053 [0.0052 0.0054] | 2123 |
| 4NT Non-Comp | 52.5 [51.8 53.2] | 0.0065 [0.0063 0.0067] | 917 |
| Flap | 68.2 [67.8 68.6] | 0.0068 [0.0067 0.0069] | 1820 |
| Hairpin | 9.6 [8.4 10.8] | 0.0034 [0.0024 0.0045] | 650 |

Given the substrate design, colocalization of donor and acceptor fluorophores should be present from the initiation of imaging. Thus, the fraction of DNA molecules exhibiting colocalization provides a direct measurement of the labeling efficiency for the different DNAs. All substrates showed comparable labeling between 64-83%, suggesting that different DNA end structures did not inhibit labeling, end annealing, or ligation (Supplementary Fig. 2A). The number of colocalized molecules from each experiment was used as the total possible number of FRET events to make accurate comparisons between the percent of FRET positive molecules over different DNA substrates. On individual colocalized molecules, we observe steady green (Figure 2B, top) and red (Figure 2B, bottom) emission upon their respective excitation, since both fluorophores are bound to the same DNA molecule. With time, individual DNAs shift to a high FRET state, observed as a decrease in green emission and a concomitant increase in red emission upon green excitation, confirming the transition from the LRC to the SRC (Figure 2B, middle at the grey dashed line).

We monitored SRC formation as the number of FRET positive molecules that accumulated over time. For blunt DNA ends, the cumulative fraction bound showed an exponential increase with 58% of molecules undergoing SRC formation at an observed rate of 0.011 s^−1^ (Figure 2D, Table 3, Supplementary Figure 2B). Interestingly, different DNA end structures (Figure 2C) showed variability in SRC formation, unlike LRC formation. Most DNA ends, 3ʹP, 5ʹOH, 1NT comp fill-in, 1NT non-comp, 4NT non-comp and flap were comparable to blunt ends with the number of molecules undergoing SRC formation ranging from 53 – 76% at observed rates from 0.005 – 0.011 s^−1^ (Figure 2D, Table 3, Supplementary Figure 2B). However, two DNAs, the 4NT complementary and hairpin substrates showed relatively fast and slow SRC formation, respectively.

4NT complementary DNA ends showed 87% SRC formation at an observed rate of 0.087 s^−1^, which corresponds to a nearly ten-fold faster rate than blunt DNA ends. Since the ends are complementary, one explanation for an increase in the number of FRET positive molecules may be self-annealing that occurs independently of the NHEJ machinery. To test this possibility, we depleted XRCC4 from HSS extract, which co-depletes its cognate binding partner, LigIV, and prevents SRC formation.^18^ In XRCC4/LigIV-depleted extract, blunt DNA ends did not form the SRC, with only 3% of DNAs undergoing FRET (Supplementary Figure 2C, top). In comparison, the 4NT complementary ends showed 90% of molecules in the high FRET state, suggesting that SRC formation is independent of NHEJ for this substrate (Supplementary Figure 2C, bottom). Furthermore, testing SRC formation in buffer alone showed 79% of molecules in the high FRET state without any extract present. Therefore, the 4NT complementary substrate appears to undergo synapsis independent of NHEJ in this assay.

In contrast, hairpin ends showed a significantly lower level of SRC formation compared to blunt ends with only 10% of ends undergoing FRET at an observed rate of 0.003 s^−1^ (Figure 2D, Table 3). We then asked whether the observed defect in SRC formation was due to a defect in LigIV recruitment. To test this, we compared DNA binding, via colocalization, and DNA end engagement, via FRET, of Cy5-labeled ybbR-LigIV/XCC4 to blunt, 1NT comp fill-in, and hairpin surface DNA substrates in buffer. All three substrates showed comparable binding amplitudes, observed rates and lifetimes for both colocalization and FRET (Supplementary Figure 2D). Since LigIV/XRCC4 binds hairpins both internally (colocalization) and at DNA ends (FRET), these data suggest that LigIV/XRCC4 binds hairpin ends normally.

The remaining substrates showed comparable SRC formation kinetics despite their distinct end chemistries and processing requirements. Thus, as we observed for the LRC, SRC formation is largely insensitive to the DNA end structure across the non-hairpin substrates tested.

### Short-range complex stability

Because the high FRET state in the SRC is indistinguishable from the ligated DNA product in our circularization assay, we sought an approach that reports SRC lifetime independent of ligation. We therefore depleted endogenous XRCC4/LigIV from HSS and restored SRC assembly by add-back of wild-type XRCC4 with catalytically dead LigIV (^cd^LigIV; LigIV K278R/XRCC4) at endogenous concentrations (Supplementary Figure 4A).^17,18,36^ Under these conditions, DNA ends can be synapsed into the SRC but cannot be covalently joined^37^, allowing the lifetime of the high-FRET state to be interpreted as SRC persistence (Supplementary Figure 3B).

SRC formation in ^cd^LigIV-rescued extract, was similar to wild-type extract (compare Figs. 2A and 3A, Tables 3 and 4), indicating that ^cd^LigIV supports the transition to the SRC with minimal perturbation. We next quantified SRC stability by measuring the duration of individual high-FRET events (rastergrams shown in Supplementary Figure 3C). For blunt ends, dwell times were best described by a double-exponential distribution with a short-lived population of 18 seconds that accounted for 73% of events and a long-lived population of 198 seconds duration that accounted for 27% of events. Previous work from our laboratory showed that LigIV-XRCC4 binds DNA ends with a dwell time of 15 seconds.^17^ This similarity suggests that the short-lived high-FRET events likely reflect an initial, LigIV-XRCC4-dependent SRC that has not yet been reinforced by additional end-bridging interactions. Subsequent formation of these additional protein-mediated contacts would convert a subset of events into the long-lived, stabilized SRC population.

**Figure 3:**
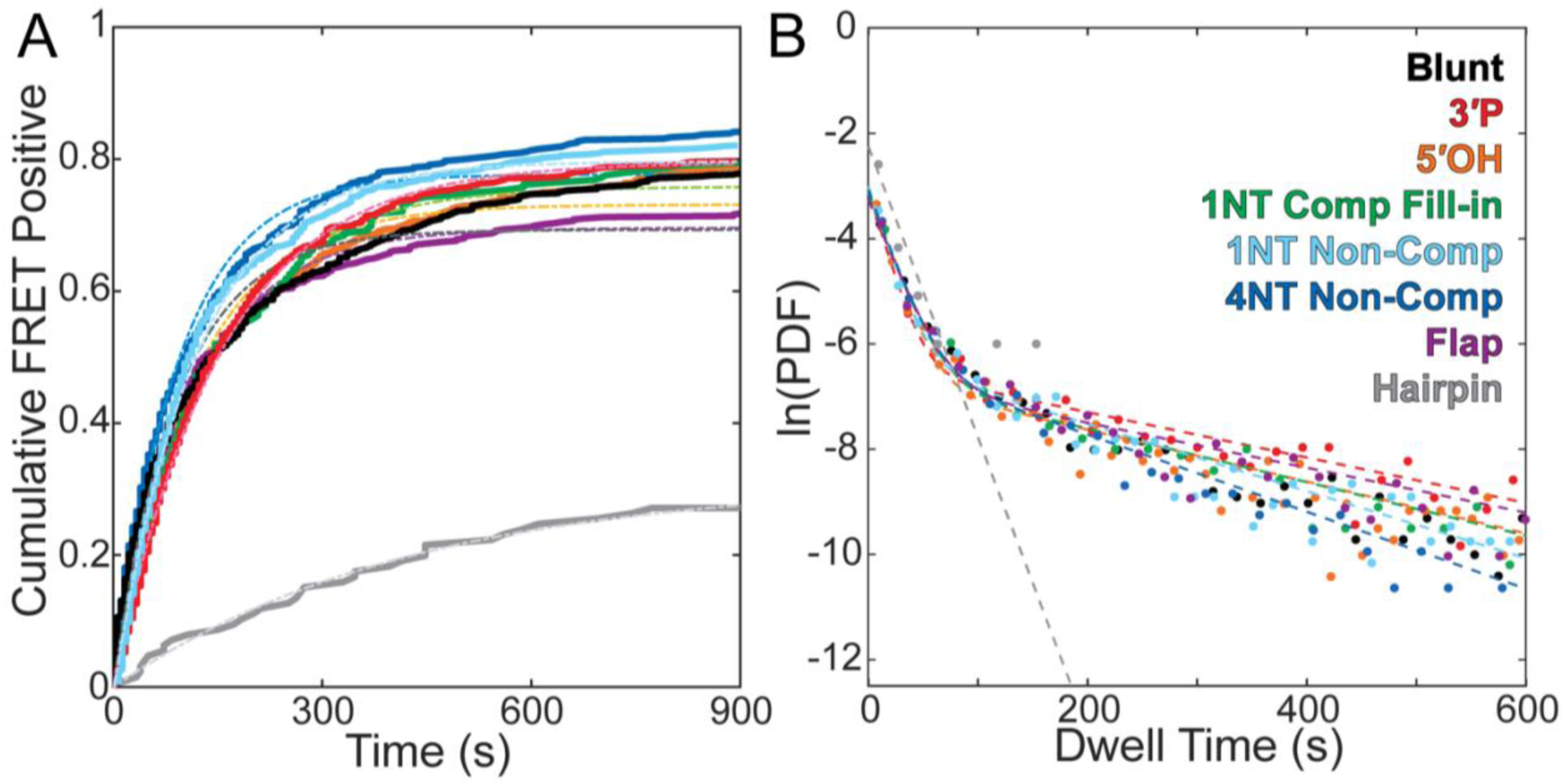
DNA end structure does not affect SRC stability. (A) SRC stability was measured in ΔXRCC4/LigIV *Xenopus* HSS extract with ^cd^LigIV/XRCC4 add-back. The cumulative fraction of FRET positive molecules, sorted by time to first FRET, shows an exponential increase over time for the different DNA substrates. Time to first binding events were fit to a single exponential model, fits shown in Table 4; rastergrams shown in Supplementary Figure 3C. (B) Dwell time histograms of individual DNA binding events were best described by a double exponential decay and plotted on a natural log scale to compare the relative stability on each DNA substrate; histograms shown in Supplementary Figure 3D, fits in Table 5.

**Table 4.** DNA Circularization SRC Formation Rates for ^cd^LigIV Extract.

| DNA | Amp (%)<br>[95% CI] | | $k_{obs}$ (s<sup>-1</sup>)<br>[95% CI] | | N |
| --- | --- | --- | --- | --- | --- |
| Blunt | 69.3 | [68.8 69.8] | 0.0108 | [0.0106 0.0110] | 1000 |
| 3'P | 80.0 | [79.3 80.7] | 0.0066 | [0.0064 0.0067] | 632 |
| 5'OH | 73.2 | [72.8 73.5] | 0.0088 | [0.0087 0.0089] | 1385 |
| 1NT Comp Fill-in | 75.9 | [75.0 76.9] | 0.0076 | [0.0073 0.0078] | 629 |
| 1NT Non-Comp | 90.5 | [90.1 90.9] | 0.0124 | [0.0122 0.0126] | 939 |
| 4NT Non-Comp | 77.7 | [77.1 78.3] | 0.0108 | [0.0105 0.0110] | 641 |
| Flap | 69.7 | [69.0 70.4] | 0.0090 | [0.0088 0.0092] | 881 |
| Hairpin | 31.3 | [26.6 36.1] | 0.0024 | [0.0018 0.0030] | 429 |

**Table 5.** Circularization DNA substrate SRC FRET Dwell Times for ^cd^LigIV Extract.

| DNA End | Amp<sub>1</sub> (%)<br>[95% CI] | $\tau_1$ (%)<br>[95% CI] | Amp<sub>2</sub> (%)<br>[95% CI] | $\tau_2$ (%)<br>[95% CI] | N |
| --- | --- | --- | --- | --- | --- |
| Blunt | 73 [70 76] | 18.0 [16.5 19.6] | 27 [24 30] | 197.9 [178.4 220.4] | 3062 |
| 3'P | 62 [59 65] | 13.8 [12.3 15.2] | 38 [35 41] | 231.1 [212.1 254.4] | 1536 |
| 5'OH | 74 [72 76] | 15.4 [14.6 16.2] | 26 [24 28] | 203.7 [189.0 219.4] | 4727 |
| 1NT Comp Fill-in | 74 [69 79] | 17.8 [15.3 20.3] | 26 [21 31] | 197.9 [166.1 235.2] | 900 |
| 1NT Non-Comp | 70 [67 73] | 14.9 [13.8 16.0] | 30 [27 33] | 156.3 [142.0 172.2] | 2889 |
| 4NT Non-Comp | 74 [70 78] | 17.0 [15.2 18.7] | 26 [22 30] | 136.8 [116.9 158.8] | 1703 |
| Flap | 70 [67 73] | 18.9 [17.2 20.7] | 30 [27 33] | 234.3 [211.4 259.7] | 1948 |
| Hairpin | 100 | 15.3 [10.5 21.3] | - | - | 75 |

Across non-hairpin substrates, SRC dwell times and population weightings were similar, with the unstable SRC making up 62 – 83% of molecules binding for 13.5 – 19 seconds and the stabilized SRC accounting for 17 – 38% of molecules bound for 137 – 234 seconds (Figure 3B, Table 5, Supplementary Figure 3D). Again, the comparable rates for each phase suggest DNA end structure is not a major determinant of SRC stability. In contrast, hairpin DNA ends lacked the long-lived population and were fit by a single ∼15 second timescale, consistent with formation of a transient SRC that fails to mature into a stabilized SRC.

Together, these data indicate that, for non-hairpin substrates, SRC stability is largely independent of DNA end chemistry. A key alternative explanation, however, is that rapid end processing in extract could convert distinct termini closer to uniform blunt ended intermediates during the measurement window. We therefore next tested whether depleting cognate processing factors reveals end-dependent effects on SRC formation or stability.

### Effect of processing factor depletion on short-range complex formation and stability

If the similarity in SRC stability across end structures were primarily driven by rapid end processing in extract, then depleting the relevant processing factors should impact the formation or stability of the SRC. To test this, we depleted individual processing factors from HSS (Supplementary Figure 4A) and measured SRC formation and stability on their cognate DNA substrates in a ^cd^LigIV-rescued background.

Depletion of Pol λ from HSS stalls end joining on a 1NT comp fill-in substrate, with Pol λ add-back recovering end joining (Supplementary Figure 4B). Previous work has confirmed this for both PNKP and Pol μ on their cognate substrates.^16^ Loss of these processing factors, however, did not substantially change the fraction of DNA molecules that formed the SRC (67 – 86%) or their rate of formation (0.009 – 0.0125 s^−1^). Thus, processing factors are not required to assemble the SRC on these damaged ends (Figure 4A, Supplementary Figure 4C).

**Figure 4:**
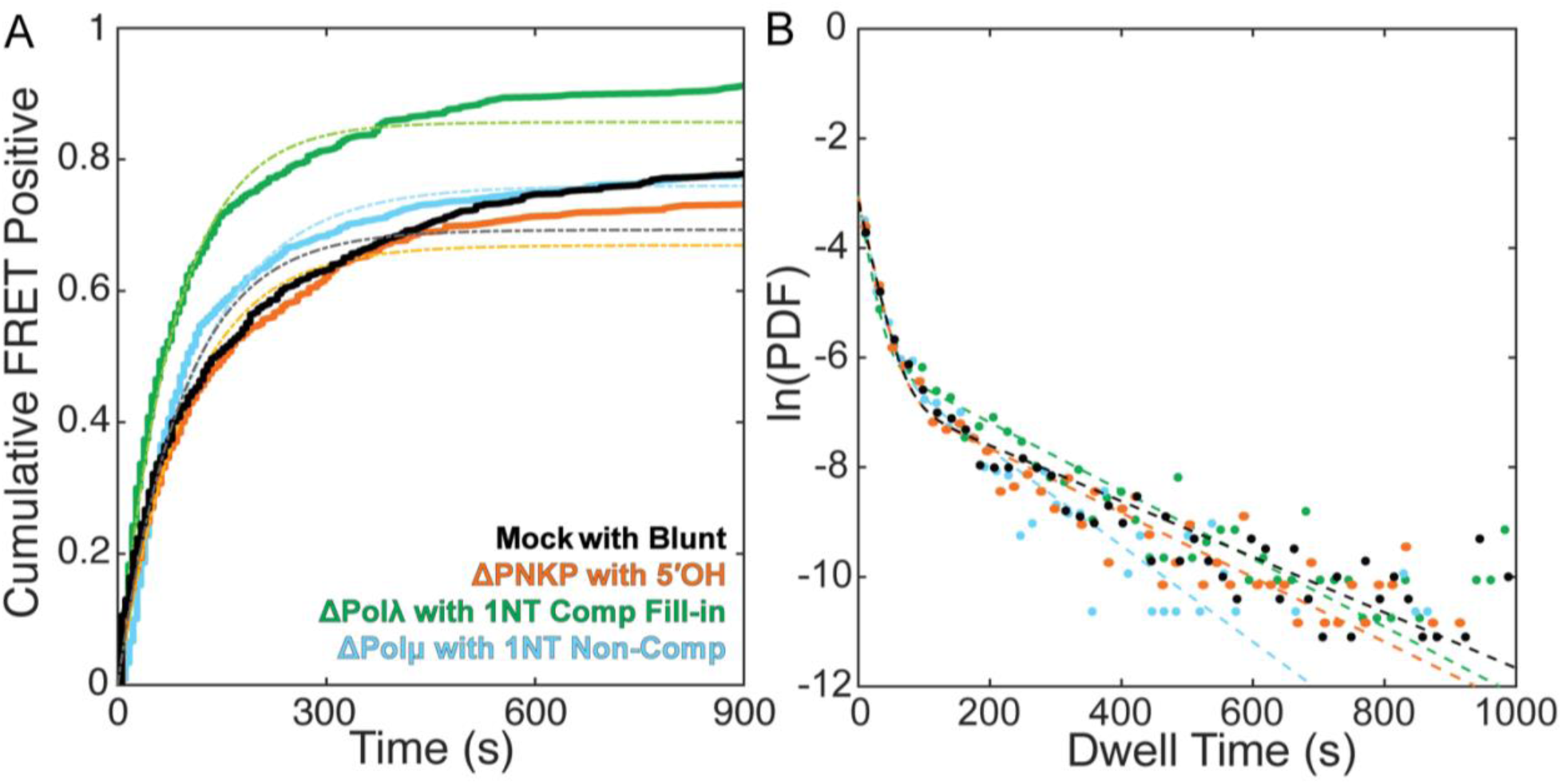
Processing factor depletion does not alter SRC formation or stability. (A) ^cd^LigIV/XRCC4 add-back in HSS extract in the background of processing factor depletion shows an increase in cumulative percent FRET positive molecules over time. Time to the first FRET event were fit to a single exponential model with fits shown in Table 6; rastergrams shown in Supplementary Figure 4B. (B) Dwell time histograms of individual DNA binding events were best described by a double exponential decay and plotted on a natural log scale to compare the relative stability on each DNA substrate; histograms shown in Supplementary Figure 4C, fits in Table 7.

**Table 6.** DNA Circularization SRC Formation Rates for ^cd^LigIV and Processing Factor Depleted Extract.

| DNA | Condition | Amp (%)<br>[95% CI] | $k_{obs}$ (s<sup>-1</sup>)<br>[95% CI] | N |
| --- | --- | --- | --- | --- |
| Blunt | cdLigIV | 69.3 [68.8 69.8] | 0.0108 [0.0106 0.0110] | 1000 |
| 5'OH | cdLigIV ΔPNKP | 67.1 [66.5 67.7] | 0.0102 [0.0101 0.0110] | 823 |
| 1NT Comp Fill-in | cdLigIV ΔPolλ | 85.8 [85.4 86.2] | 0.0125 [0.0123 0.0126] | 789 |
| 1NT Non-Comp | cdLigIV ΔPolμ | 75.9 [75.3 76.5] | 0.009 [0.0088 0.0091] | 849 |

**Table 7.** Circularization DNA substrate SRC FRET Dwell Times for ^cd^LigIV and Processing Factor Depleted Extract.

| DNA End | Condition | Amp<sub>1</sub> (%)<br>[95% CI] | $\tau_1$ (%)<br>[95% CI] | Amp<sub>2</sub> (%)<br>[95% CI] | $\tau_2$ (%)<br>[95% CI] | N |
| --- | --- | --- | --- | --- | --- | --- |
| Blunt | cdLigIV | 73 [70 76] | 18.0 [16.5 19.6] | 27 [24 30] | 198 [178 220] | 3062 |
| 5'OH | cdLigIV ΔPNKP | 75 [71 77] | 16.9 [15.8 18.2] | 25 [23 29] | 168 [150 189] | 2533 |
| 1NT Comp Fill-in | cdLigIV ΔPolλ | 58 [54 61] | 12.8 [11.7 14.0] | 42 [39 46] | 161 [147 176] | 2161 |
| 1NT Non-Comp | cdLigIV ΔPolμ | 70 [66 74] | 16.2 [14.7 17.8] | 30 [26 34] | 115 [102 129] | 2298 |

We next asked whether these processing factors contribute to maintaining SRCs once formed. SRC dwell time distributions were well described by short- and long-lived populations with parameters similar to undepleted conditions (Figure 4B, Table 7, Supplementary Figure 4D). Although the long-lived population was modestly reduced in some depletions, the overall stability of the SRC was retained, demonstrating that processing factors do not have a significant role in establishing or maintaining synapsis. Together, these results argue against a model in which rapid processing “erases” DNA end differences. Instead, they support a model in which core NHEJ proteins create a protein scaffold that maintains SRC synapsis across diverse end chemistries, while processing factors act within this synaptic complex.

### XLF tail-Ku interactions promote SRC formation on ends requiring processing

The SRC consists of a stabilized network of protein-protein interactions that bridge DNA ends. A central player in this network is XLF. XLF engages XRCC4 though head-to-head interactions that are required for SRC formation, while its C-terminal tails contain Ku-binding motifs (KBMs) that can contact Ku at DNA ends.^35,38–41^ In addition, cryo-EM structures identified contacts between the XLF stalk and Ku, although the functional contribution of these interactions remains unclear.^42^ Thus, XLF is positioned to coordinate multiple contacts across the break rather than acting through a single protein bridge.

Previous work from our laboratory showed that an asymmetric XLF heterodimer containing one wild-type subunit and one subunit lacking the C-terminal tail and KBM supports efficient blunt-end joining.^35^ This one intact XLF KBM-Ku interaction is sufficient when DNA ends can be ligated without processing. We reasoned that this single interaction may not be sufficient on substrates that cannot be ligated immediately. In this setting, two XLF KBM-Ku interactions may promote formation or maintenance of the SRC while processing factors access the DNA ends.

To test this possibility, we compared a ^WT/WT^XLF heterodimer, which contains two intact tails and KBMs, with a ^WT/ΔTailΔKBM^XLF heterodimer, which retains the essential XLF-XRCC4 head interaction and one intact tail/KBM but lacks the second tail/KBM-mediated Ku interaction. Using these XLF heterodimer constructs, we first tested bulk end-joining on a blunt-ended DNA substrate to confirm comparable joining efficiency using a radioactive gel-based assay.^35^ In wild-type HSS extract, the linear input DNA was first converted to the singly ligated open-circular product and then to the doubly ligated supercoiled product over the time course (Figure 5A, lanes 1-5). Consistent with prior work, XLF depletion abolished end-joining while add-back of either the ^WT/WT^XLF or ^WT/ΔTailΔKBM^XLF heterodimer showed a robust recovery of end-joining (Figure 5A). Quantification of the products revealed only a modest difference in joining between the two XLF constructs (Figure 5C).

**Figure 5:**
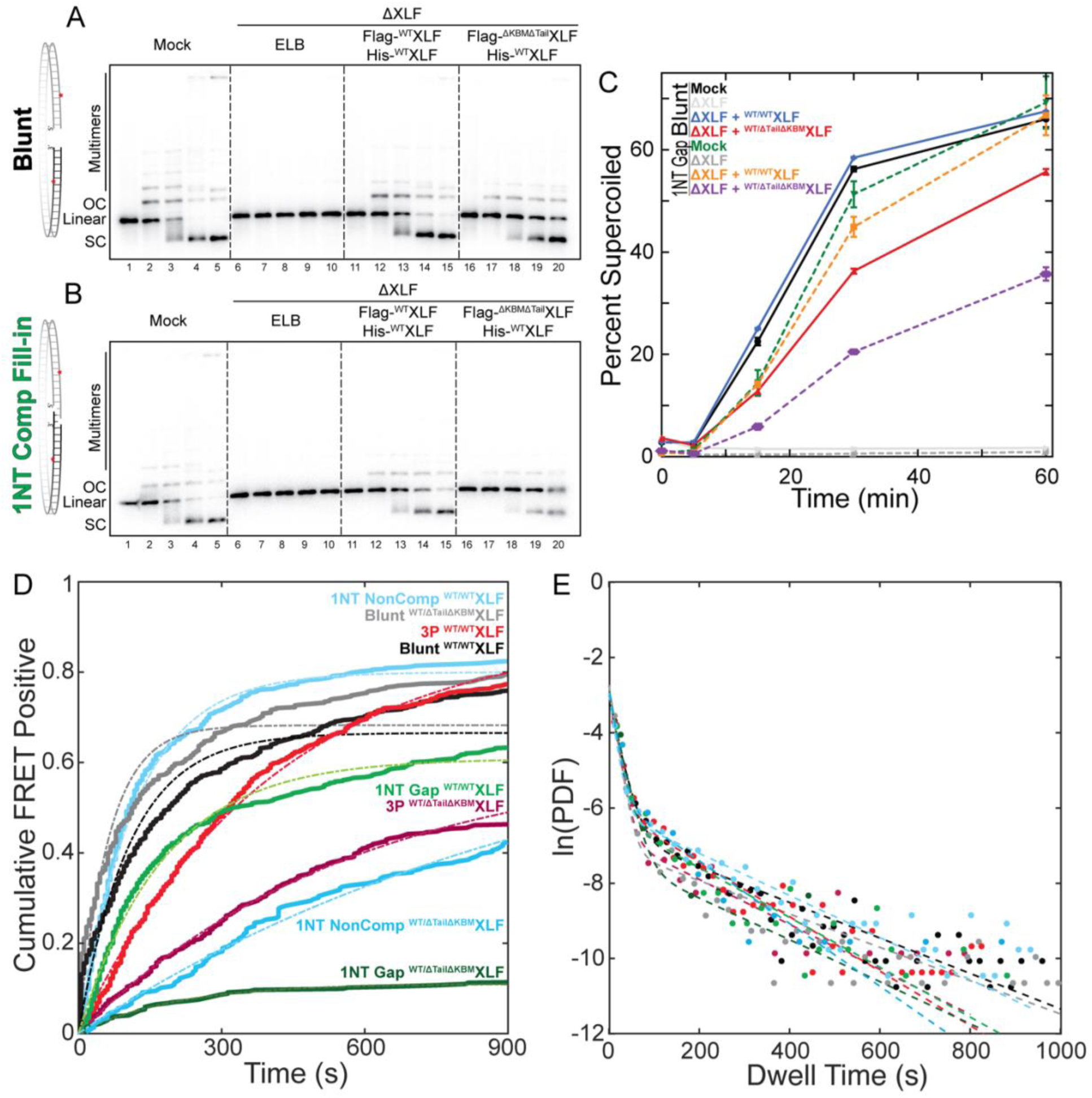
XLF promotes SRC formation on DNA ends requiring end processing. Bulk end joining assay comparing the activity of different XLF heterodimers to mock depleted extract for joining blunt (A) and 1NT complementary fill-in (B) DNA substrates, quantified in (C). (D) XLF heterodimer add-back in HSS extract in the background of ^cd^LigIV/XRCC4 shows an increase in cumulative percent FRET positive molecules over time. Molecules sorted by the time to the first FRET event were fit to a single exponential model with fits shown in Table 8; rastergrams shown in Supplementary Figure 5A. (E) Dwell time histograms of individual DNA binding events were best described by a double exponential decay and plotted on a natural log scale to compare the relative stability on each DNA substrate; histograms shown in Supplementary Figure 5B, fits in Table 9.

**Table 8.** DNA Circularization SRC Formation Rates for ^cd^LigIV and XLF Depleted Extract.

| DNA End | Condition | Amp (%)<br>[95% CI] | $k_{obs}$ (s<sup>-1</sup>)<br>[95% CI] | N |
| --- | --- | --- | --- | --- |
| Blunt | cdLigIV / <sup>WT/WT</sup>XLF | 66.6 [66.0 67.1] | 0.0093 [0.0091 0.0095] | 684 |
| Blunt | cdLigIV / <sup>WT/ΔTailΔKBM</sup>XLF | 68.3 [67.8 68.8] | 0.0159 [0.0155 0.0163] | 625 |
| 3'P | cdLigIV / <sup>WT/WT</sup>XLF | 89.3 [88.0 90.6] | 0.0025 [0.0024 0.0026] | 654 |
| 3'P | cdLigIV / <sup>WT/ΔTailΔKBM</sup>XLF | 61.7 [58.1 65.4] | 0.0018 [0.0016 0.0019] | 503 |
| 1NT Comp Fill-in | cdLigIV / <sup>WT/WT</sup>XLF | 60.9 [60.3 61.6] | 0.0057 [0.0055 0.0058] | 732 |
| 1NT Comp Fill-in | cdLigIV / <sup>WT/ΔTailΔKBM</sup>XLF | 11.0 [9.6 12.5] | 0.0050 [0.0038 0.0062] | 492 |
| 1NT Non-Comp | cdLigIV / <sup>WT/WT</sup>XLF | 80.0 [79.3 80.6] | 0.0084 [0.0082 0.0086] | 565 |
| 1NT Non-Comp | cdLigIV / <sup>WT/ΔTailΔKBM</sup>XLF | 74.5 [65.4 83.5] | 0.0010 [0.0008 0.0011] | 390 |

**Table 9.**
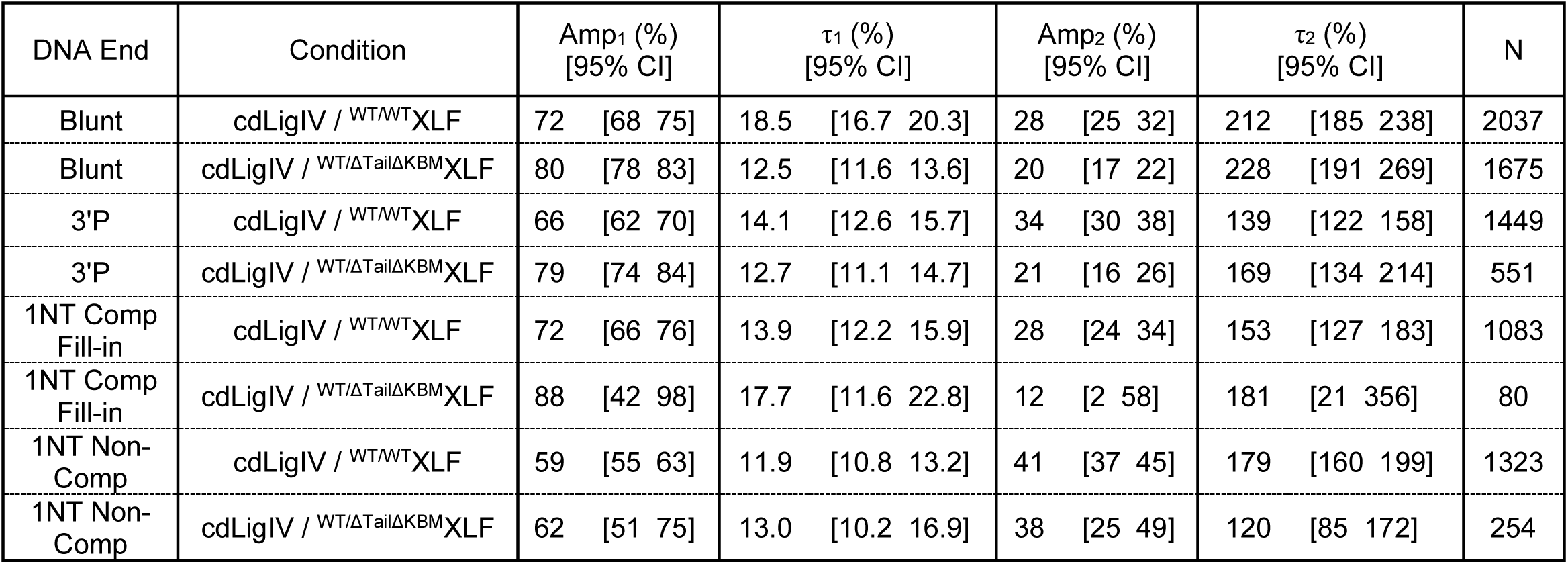
Circularization DNA substrate SRC FRET Dwell Times for ^cd^LigIV and XLF Depleted Extract.

| DNA End | Condition | Amp<sub>1</sub> (%)<br>[95% CI] | $\tau_1$ (%)<br>[95% CI] | Amp<sub>2</sub> (%)<br>[95% CI] | $\tau_2$ (%)<br>[95% CI] | N |
| --- | --- | --- | --- | --- | --- | --- |
| Blunt | cdLigIV / <sup>WT/WT</sup>XLF | 72 [68 75] | 18.5 [16.7 20.3] | 28 [25 32] | 212 [185 238] | 2037 |
| Blunt | cdLigIV / <sup>WT/ΔTailΔKBM</sup>XLF | 80 [78 83] | 12.5 [11.6 13.6] | 20 [17 22] | 228 [191 269] | 1675 |
| 3'P | cdLigIV / <sup>WT/WT</sup>XLF | 66 [62 70] | 14.1 [12.6 15.7] | 34 [30 38] | 139 [122 158] | 1449 |
| 3'P | cdLigIV / <sup>WT/ΔTailΔKBM</sup>XLF | 79 [74 84] | 12.7 [11.1 14.7] | 21 [16 26] | 169 [134 214] | 551 |
| 1NT Comp Fill-in | cdLigIV / <sup>WT/WT</sup>XLF | 72 [66 76] | 13.9 [12.2 15.9] | 28 [24 34] | 153 [127 183] | 1083 |
| 1NT Comp Fill-in | cdLigIV / <sup>WT/ΔTailΔKBM</sup>XLF | 88 [42 98] | 17.7 [11.6 22.8] | 12 [2 58] | 181 [21 356] | 80 |
| 1NT Non-Comp | cdLigIV / <sup>WT/WT</sup>XLF | 59 [55 63] | 11.9 [10.8 13.2] | 41 [37 45] | 179 [160 199] | 1323 |
| 1NT Non-Comp | cdLigIV / <sup>WT/ΔTailΔKBM</sup>XLF | 62 [51 75] | 13.0 [10.2 16.9] | 38 [25 49] | 120 [85 172] | 254 |

We next asked whether the XLF heterodimers could rescue joining of a 1NT comp fill-in substrate that requires processing before joining. Wild-type extract and ^WT/WT^XLF heterodimer add-back showed comparable repair to the blunt end condition, while XLF depletion again eliminated joining (Figure 5B, lanes 1-15). In contrast, the ^WT/ΔTailΔKBM^XLF heterodimer restored joining of the 1NT complementary fill-in substrate substantially less efficiently than ^WT/WT^XLF (Figure 5B, lanes 16-20; Figure 5C). These findings suggest that the second XLF tail/KBM-mediated Ku interaction becomes functionally important when ligation is delayed by a requirement for end processing.

The reduced joining of processing-dependent ends could result from defective SRC formation and/or reduced stability of the SRC. We therefore used the ^cd^LigIV-background smFRET assay to investigate these possibilities. On blunt DNA ends, both the ^WT/WT^XLF and ^WT/ΔTailΔKBM^XLF heterodimers supported comparable SRC formation, with 67 – 68% of molecules undergoing FRET and similar observed rates (*k_obs_* = 0.01 – 0.016 s^−1^), comparable to wild-type extract (Figure 5D, Table 8, Supplementary Figure 5A). We next examined three end structures that require end processing prior to ligation, a 3ʹP end (processed by PNKP), a 1NT complementary end requiring fill-in (processed by Polλ), and a 1NT non-complementary end (processed by Polμ). With ^WT/WT^XLF heterodimer, all three substrates showed comparable SRC formation to blunt DNAs (61 – 89 % FRET positive, *k_obs_* = 0.002 – 0.005 s^−1^; Figure 5D, Table 8). In contrast, the ^WT/ΔTailΔKBM^XLF heterodimer showed reduced SRC formation on all three processed DNA substrates, with a 50% decrease or greater in FRET positive molecules over 900 seconds (*k_obs_* = 0.001 – 0.005 s^−1^; Figure 5D, Table 8). Because the apparent rates among complexes that did form were similar, the primary defect is a reduced probability of productive SRC formation rather than a uniform slowing of SRC assembly.

The SRCs that formed with ^WT/ΔTailΔKBM^XLF displayed near normal dwell time distributions (transient population: 59 – 88 %, dwell time = 12.5 – 18.5 s; stable population: 12 – 41 %, dwell time = 120 – 228 s). Thus, the second XLF tail/KBM-mediated Ku interaction appears to act primarily during the establishment of a productive SRC on processing-dependent ends. Other contacts within the NHEJ bridging network, including the remaining XLF-Ku interactions, and by PAXX, may stabilize the subset of SRCs that form.

## Discussion

NHEJ must retain the correct pair of DNA ends while repairing termini that are often incompatible with immediate ligation.^43^ Using physiologically complex *Xenopus* egg extract that carries out robust end processing and joining, we systematically investigated how a diverse panel of DNA end chemistries requiring PNKP, polymerase or nuclease processing impacts the formation and stability of the LRC and SRC. We show that despite vast differences in the chemical nature of these substrates, formation of the LRC and subsequent transition to the SRC are largely independent of the DNA ends. Hairpin-containing ends are an important exception since they readily form the LRC but are significantly delayed in transitioning to the SRC, and 4NT complementary ends appear to synapse spontaneously at high concentrations independent of the NHEJ machinery. Our results add to a growing understanding that DNA end synapsis is mediated by a highly redundant network of protein-protein interactions that enable synapsis, processing, and joining.^4^ However, as we demonstrate for XLF, some interactions become increasingly important when processing is required.

The similar LRC kinetics observed across our substrate panel are consistent with LRC assembly being dominated by interactions between DNA-PK complexes rather than direct recognition of DNA ends examined here.^26,42^ Within the LRC, DNA ends are held apart by the DNA-PKcs dimer interface, with PAXX providing an additional bridge between Ku molecules on opposing ends.^23,42^ Such an architecture provides a straightforward explanation for why chemical modifications, short overhangs and flapped substrates do not alter LRC formation or lifetime. This conclusion does not imply that DNA ends make no contribution to DNA-PK function. Rather, it suggests that chemical diversity represented by the substrates we examined does not strongly influence the initial protein-mediated synapsis step.

Hairpin termini were a notable exception. Hairpin substrates readily formed LRCs but showed strongly reduced SRC formation and lacked the long-lived SRC population observed with other substrates. Hairpins are intermediates in V(D)J recombination and engage DNA-PK differently from duplex DNA ends.^44^ Biochemical and structural work has shown that hairpin- and duplex-bound DNA-PKcs adopt distinct conformations and phosphorylation states, in part because of differences in how the DNA-PKcs end-capping helix engages the DNA terminus.^44^ Our observation that LigIV/XRCC4 binds comparably to hairpin and blunt substrates argues that impaired LigIV/XRCC4 recruitment alone is unlikely to explain the observed defect in SRC formation for hairpin ends. Thus, this defect is consistent with the possibility that hairpin-bound DNA-PK adopts an architecture less permissive for SRC formation.^44^ However, our experiments do not directly identify the responsible DNA-PK conformation or distinguish other possibilities including the effects of Artemis-dependent processing or other steps in the LRC-to-SRC transition.

A potential explanation for the similar SRC kinetics observed across chemically distinct ends is that processing factors in extract rapidly convert diverse termini into a common intermediate before SRC formation and stability are measured. Our processing factor depletion experiments, however, argue against this explanation. Depletion of PNKP, Pol λ or Pol μ impaired repair of their cognate substrates in bulk end joining reactions, yet did not substantially alter SRC formation or stability in the ^cd^LigIV-rescued single-molecule assay. Although some depletion conditions modestly affected the long-lived SRC population, stable synapsis remained. Thus, for the processing reactions examined here, the cognate processing enzymes are not major structural determinants of SRC assembly or maintenance. Instead, these results are consistent with processing reactions within an SRC that is principally maintained by core NHEJ factors. Whether this remains true on DNA with more complex lesions such as bulky adducts or chromatin-containing templates remains to be determined.

Our XLF-tail results identify a condition in which the apparent redundancy of the SRC bridging network is reduced. Previous work from our laboratory showed that a single LigIV molecule bridges DNA ends in the SRC and promotes an initial ligation attempt when ends are compatible.^17^ On ends that cannot be ligated immediately, this configuration must be remodeled to permit processing factor access, either through LigIV dissociation or conformational opening, as suggested by a recent cryo-EM structure of a Polμ−containing SRC.^27^ We propose that this interval creates a greater requirement for additional bridging interactions to establish or preserve a productive synaptic complex. Consistent with this model, an XLF heterodimer retaining only one intact XLF tail/KBM supported blunt-end joining and SRC formation but impaired both joining and SRC formation on a Polλ−dependent fill-in substrate. It also reduced productive SRC formation on PNKP- and Polμ−dependent substrates, suggesting that a second XLF tail/KBM interaction becomes important across distinct end-processing contexts. The near-normal dwell times of the SRCs that did form suggest that loss of one XLF tail/KBM primarily reduces the probability of establishing a productive SRC, rather than destabilizing all SRCs after their formation. Although we have not systematically tested every protein-protein interaction within the SRC, our findings suggest that other contacts that are redundant during repair of ligation-competent ends may become functionally important when end processing delays ligation.

We propose that NHEJ synapsis is maintained by a network of partially redundant protein-mediated bridges (Figure 6).^29,42,43,45–51^ Across the non-hairpin end structures examined here, this network supports formation and persistence of the LRC and SRC despite distinct DNA end chemistries and processing requirements. When ligation is delayed, XLF KBM-mediated interactions with Ku promote the formation of productive SRCs, consistent with processing occurring within the SRC. In this model, DNA end chemistry is not primarily sensed at the level of synaptic complex assembly. Instead, it impacts the selection and execution of downstream processing reactions that determine repair outcome.

**Figure 6:**
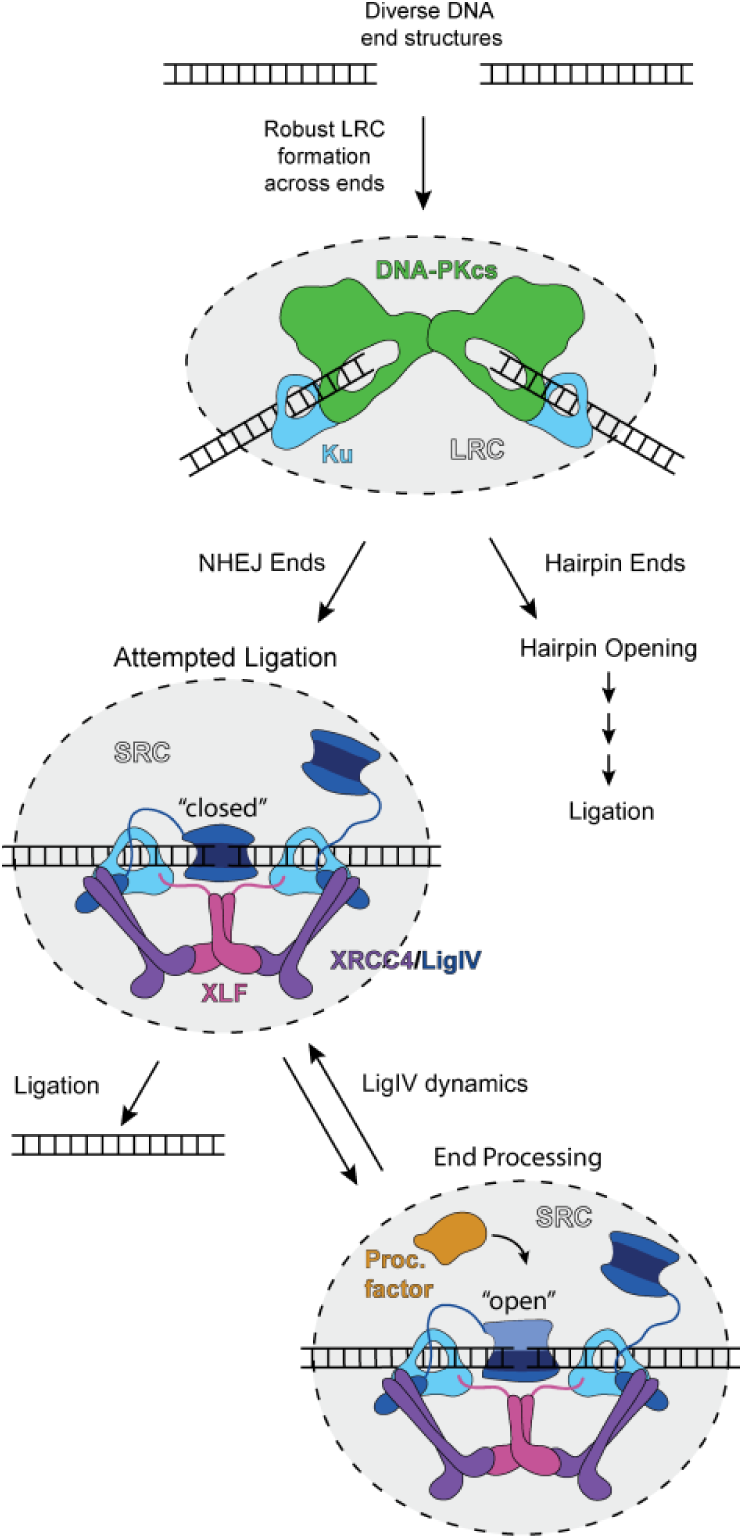
Model of how NHEJ accommodates diverse DNA end structures. After initial formation of the LRC, hairpin ends are processed in a distinct manner from other canonical NHEJ ends. Once NHEJ competent broken ends are tethered inside the SRC, protein-protein interactions between core NHEJ factors maintain the SRC to keep the ends in close proximity during iterative rounds of processing and attempted ligation. Conformational opening of LigIV (depicted), or its dissociation, allows processing factors access to the DNA ends to enable repair.

## Methods

### Xenopus egg extract preparation

The high-speed supernatant (HSS) of *Xenopus laevis* eggs was prepared as previously described.^52^

### Antibody production, extract immunodepletion and immunoblotting

Rabbit polyclonal antibodies against XRCC4, XLF, PNKP, Polλ, and Polμ were prepared previously.^16,18^ Histone H3 antibody was purchased from Cell Signaling. Factors were immunodepleted from HSS using 1 volume of Protein A Sepharose beads (Cytiva) to 3 volumes of purified antibody (1 mg/mL; Pocono Rabbit Farm & Laboratory) to 5 volumes of HSS extract over three rounds of depletion. Protein A Sepharose beads were removed from storage buffer and incubated with the purified antibody on a rotating wheel at 4°C overnight. Beads were then washed through four rounds with a ten-fold excess of ELB (10 mM HEPES, pH 7.7, 50 mM KCl, 2.5 mM MgCl_2_, 250 mM sucrose). HSS extract was supplemented with 7.5 ng/μL nocodazole before addition to the washed beads and rotated for 1 hour at 4°C. After 1 hour, extract was moved to fresh beads and repeated a total of three times. Depleted extract was aliquoted and flash frozen in liquid nitrogen before storing at −80°C until needed.

Depletions were confirmed by immunoblotting. Samples were diluted 4-fold in Laemmli buffer, separated on a 4-15% Mini-PROTEAN TGX precast gel (Bio-Rad; 32 minutes at 200 V) in SDS running buffer (Boston BioProducts) and transferred to a PVDF membrane (Perkin Elmer; 70 minutes at 300 mA) in transfer buffer (Boston BioProducts). Membranes were then blocked at room temperature with 5% nonfat milk in PBST for 75 minutes, rinsed with PBST five times and incubated overnight with a 1:3000 dilution of primary antibody in PBST containing 0.5% sodium azide. After three washes with 5% nonfat milk in PBST, blots were incubated for one hour at room temperature with a secondary goat anti-rabbit horseradish peroxidase-conjugated antibody diluted 1:20,000 in 5% nonfat milk in PBST. Blots were rinsed with PBST before imaging using WesternBright ECL-Spray (Advansta Corporation) on an Amersham Imager 6000 (GE Healthcare). Image contrast levels were adjusted for visualization as needed.

### LigIV/XRCC4 purification and labeling

Plasmids for K278R-LigIV/XRCC4 and ybbR-LigIV/XRCC4 were obtained from previous work.^17,18^ Individual plasmids were transformed into BL21 (DE3) Rosetta pLysS cells by incubating 100 ng plasmid with 50 μL of cells on ice for 1 hour before heat-shocking for 40 seconds at 42°C. After cooling on ice for 2 minutes, cells recovered in 100 μL SOC media while shaking at 37°C. After 1 hour, cells were plated on LB-ampicillin plates and left to grow overnight at 37°C. Plates were removed the next morning and then a single colony was inoculated into 10 mL Luria Broth (LB) (VWR) with 100 μg/μL ampicillin and grown overnight shaking at 37°C. The next day, 250 mL TB-Amp cultures were inoculated with 1 mL of overnight culture and allowed to grow at 37°C, shaking at 200 RPM, to an OD_600_ of 1.2 before inducing overnight with 1 mM isopropyl β-D-1-thiogalactopyranoside (IPTG) at 16°C. Cells were harvested by centrifugation at 4,667 × *g* for 5 min before resuspending in lysis buffer (20 mM Tris-HCl, pH 8, 400 mM NaCl, 10 mM imidazole, 1 mM DTT, 10% glycerol) and flash freezing in liquid nitrogen for storage at −80°C.

Resuspended cells were thawed and supplemented with a cOmplete, ethylenediaminetetraacetic acid (EDTA) free protease inhibitor cocktail tablet (Roche) pre-dissolved in 1 mL of deionized water. Cells were lysed on ice by sonicating for 2.5 min active, 1 second on, 3 seconds off, at 40% amplitude (FisherBrand Sonic Dismembrator) before separating the cellular debris via centrifugation at 47,850 × *g* for 1 hour. The clarified supernatant was then incubated with a 1 mL bed volume of Ni-NTA (Qiagen) beads, pre-washed with lysis buffer, gently rotating at 4°C for 1 hour. Resin was then pelleted by centrifugation at 1,550 × *g* for 2 minutes to remove the supernatant before washing with 10 mL lysis buffer twice. Once washed, the resin was resuspended in 10 mL of lysis buffer to transfer to a 5 mL disposable column (Qiagen) and allowed to settle. Protein was eluted with 10 mL lysis buffer supplemented with 250 mM imidazole.

Protein was then applied to a 2 mL Strep-Tactin XT column (IBA LifeSciences) pre-washed with lysis buffer. The column was then washed with 10 mL of lysis buffer twice before eluting with 10 mL BXT buffer (IBA LifeSciences). Eluted protein was dialyzed overnight into Size Exclusion Chromatography (SEC) buffer (2 x 1L; 20 mM HEPES, pH 7.5, 150 mM NaCl, 10% glycerol, 1 mM DTT) before concentrating and loading onto a Superdex 200 Increase 10/300 GL column (Cytiva) run on an ÄKTA Pure (Cytiva). Pooled fractions containing pure protein were concentrated for storage before flash freezing in liquid nitrogen and storing at −80°C.

For ybbR labeling, protein eluted from the Strep-Tactin column was supplemented with 10% glycerol and 10 mM MgCl_2_ before concentration. The labeling reaction consisted of the protein, 25-fold excess Cy5-CoA, and 10 μM Sfp synthase. After rotating overnight at 4°C, the protein was run over SEC as the unlabeled version. Final protein purity shown in Supplementary Figure 3A.

### XLF purification

Plasmids for ^WT/WT^XLF and ^WT/ΔTailΔKBM^XLF heterodimers were obtained from previous work.^38^ Individual plasmids were transformed into BL21 (DE3) Rosetta pLysS cells by incubating 100 ng plasmid with 50 μL of cells on ice for 1 hour before heat-shocking for 40 seconds at 42°C. After cooling on ice for 2 minutes, cells recovered in 100 μL SOC media while shaking at 37°C. After 1 hour, cells were plated on LB-ampicillin plates and left to grow overnight at 37°C. Plates were removed the next morning and then a single colony was inoculated into 10 mL Luria Broth (LB) (VWR) with 100 μg/μL ampicillin and grown overnight shaking at 37°C. The next day, 2 × 1L LB-Amp cultures were inoculated with 3 mL of overnight culture and allowed to grow at 37°C, shaking at 200 RPM, to an OD_600_ of 0.8 before inducing overnight with 1 mM isopropyl β-D-1-thiogalactopyranoside (IPTG) at 16°C. Cells were harvested by centrifugation at 4,667 × *g* for 5 min before resuspending in lysis buffer (20 mM Tris-HCl, pH 8, 1 M NaCl, 30 mM imidazole, 5 mM β-mercaptoethanol, 10% glycerol) and flash freezing in liquid nitrogen for storage at −80°C.

Resuspended cells were thawed and supplemented with a cOmplete, ethylenediaminetetraacetic acid (EDTA) free protease inhibitor cocktail tablet (Roche) pre-dissolved in 1 mL of deionized water. Cells were lysed on ice by sonicating for 2.5 min active, 1 second on, 3 seconds off, at 40% amplitude (FisherBrand Sonic Dismembrator) before separating the cellular debris via centrifugation at 47,850 × *g* for 1 hour. The clarified supernatant was incubated with a 2 mL bed volume of Ni-NTA (Qiagen) beads, pre-washed with lysis buffer, gently rotating at 4°C for 1 hour. Resin was then pelleted by centrifugation at 1,550 × *g* for 2 minutes to remove the supernatant before washing with 30 mL lysis buffer twice. Once washed, the resin was resuspended in 10 mL of lysis buffer to transfer to a 5 mL disposable column (Qiagen) and allowed to settle. Protein was eluted using 10 mL lysis buffer supplemented with 250 mM imidazole and concentrated to 0.5 mL using a 10 kDa MWCO Amicon Ultra concentrator (Millipore Sigma). To reduce salt and imidazole concentration, protein was then diluted with 5 mL of FLAG wash buffer (20 mM Tris-HCl, pH 8, 400 mM NaCl, 10% glycerol), reconcentrated and diluted for a total of three rounds.

Concentrated protein was then applied to 0.25 mL ANTI-FLAG M2 Affinity Gel (Sigma Aldrich) resin pre-washed with FLAG wash buffer and gently rotated at 4°C for 1 hour. Resin slurry was then transferred to a 1 mL column and washed three times with 5 mL of FLAG wash buffer. Protein was eluted with 1 mL of FLAG wash buffer supplemented with 0.5 mg/mL 3x FLAG peptide. Eluted protein was loaded onto a Superdex 200 Increase 10/300 GL column (Cytiva) run on an ÄKTA Pure (Cytiva) in Size Exclusion Chromatography (SEC) buffer (20 mM Tris-HCl, pH 8, 400 mM NaCl, 10% glycerol, 5 mM β-mercaptoethanol). Pooled fractions containing pure protein were then concentrated for storage, if needed, before flash freezing in liquid nitrogen and storing at −80°C. Final protein purity shown in Supplementary Figure 3A.

### DNA substrate production

DNA oligonucleotides were purchased from Integrated DNA Technologies; sequences are listed in Supplementary Table 1 and 2. Cy3B- or ATTO 647N-NHS Ester (Lumiprobe) labeling of amino-modified oligos was performed overnight, rotating in the dark at room temperature in 100 mM sodium tetraborate, pH 8.5, 10% dimethyl sulfoxide (DMSO) at 1 mg/mL of oligo and 2 mg/mL NHS ester. After overnight incubation, unlabeled and labeled DNAs were separated via 20% acrylamide (19:1) 7 M Urea-PAGE at 18 W for 1 hour in 0.8 x TTE glycerol tolerant buffer. The labeled DNA band was excised using UV shadowing before electro-eluting for 30 min using 3k MWCO dialysis tubing in 1x TBE buffer followed by ethanol precipitation and resuspension in 10 mM Tris–HCl, pH 8.

Radiolabeling of substrates was achieved by PNK (NEB) phosphorylation using γ-^32^P-ATP (Perkin Elmer) in PNK buffer (NEB) following the manufacturer’s protocol.

To achieve various end structures, DNA oligonucleotides were phosphorylated using PNK (NEB) following the manufacturer’s protocol, when needed, before annealing. Strands for the 3′P and 5′OH substrates were purchased from IDT and not subjected to PNK reaction. Fluorescently labeled oligo duplexes were annealed by heating to 65°C and cooling by 1°C per minute to 20°C; unlabeled duplexes were annealed by heating to 95°C and cooling by 1°C per minute to 20°C. Annealed duplexes contained sticky ends for either SphI or AatII for ligation to backbones.

LRC backbones were annealed as above. LRC substrates were ligated together at a 1:1.25 ratio of oligo duplex:backbone. After ligation, in-solution DNAs were incubated with a 2.5-fold molar excess of SavPhire monomeric streptavidin (Sigma Aldrich) for one hour before use.

SRC backbones were generated as previously described.^16^ Briefly, the plasmid backbone was digested using SphI and AatII to generate two different sticky ends as well as *Nt*.BbvCI to anneal in a biotinylated strand.

The backbone was then gel-separated and purified before annealing both oligo duplexes and the biotinylated strand in 1:10:10:10 ratio in the presence of T4 DNA ligase (NEB) in T4 DNA ligase buffer. Ligation was carried out at 16°C for one hour and then run out on a 1% agarose gel in TBE. Linear DNA molecules were electroeluted in 3.5 kDa MWCO SnakeSkin dialysis tubing (ThermoFisher), ethanol precipitated, and resuspended in 10 mM Tris-HCl, pH 8.

### Bulk assays

End-joining by *Xenopus* HSS egg extract was monitored by gel shift of the radiolabeled DNA upon separation on an agarose gel. Reactions consisted of mock or depleted HSS extract supplemented with 3 ng/μL radiolabeled DNA substrate, 5 ng/μL carrier DNA, ATP regeneration system (ARS; 1.5 mM ATP, 7.5 mM Phosphocreatine, 0.01 mg/mL creatine phosphokinase) and 50 nM XLF heterodimer. At the indicated time points, 2 μL of the reaction was quenched in 5 μL replication stop buffer (80 mM Tris-HCl, pH 8, 8 mM EDTA, 10% Ficoll, 5% SDS, 0.13% phosphoric acid, 0.2% bromophenol blue) before adding proteinase K (1.5 mg/mL final) for 1 hour at 37°C. Samples were separated on a 1% agarose gel run for 1.5 hours. Gels were dried on Hybond (Cytiva) and visualized using a Typhoon FLA 7000 (GE Healthcare). Gel bands were quantified using Multi Gauge (FujiFilm) and plotted as the percent intensity of the supercoiled band compared to total lane intensity. Error bars represent the standard error of the mean from a biological duplicate.

### Single-molecule assay set-up

Single-molecule microscope set-up and flow cell assembly were the same as previously described^16,17,53^. Surface laser power density was 750 mW/cm^2^ for the 532 nm laser and 500 mW/cm^2^ for the 642 nm laser, as measured on the table using a Coherent FieldMate power meter. Flow cells were made in the same manner as previously described.^17^

Coverslips were prepared by washing number 1.0 glass coverslips with ethanol, water, 1 M KOH, and water for 30 minutes each, sonicating; this was repeated twice. Coverslips were then coated with 2% 3-aminopropyltriethoxysilane and cured for 45 minutes in a drying oven at 100°C. After cooling, coverslips were sandwiched together in a PEGylating solution with a 10:1 ratio of 4-arm PEG:Biotin PEG SVA (Laysan Bio) in 100 mM sodium bicarbonate before washing with deionized water and storing at −20°C.

Flow cells were constructed from glass slides with holes drilled 10 mm apart to allow attachment of PE20 and PE60 tubing (BD Intramedic) by epoxy (DevCon). A 1.5 x 12 mm channel was cut from a double-sided SecureSeal Adhesive Sheet (Grace Bio-Labs) and fixed to the other face of the glass slide. The functionalized coverslip was then stuck to the slide and sealed with epoxy. Solutions were introduced into the flow cell using a 1 mL syringe connected to the PE60 tubing with a 21-gauge needle.

### Two-piece DNA binding assay

The two-piece DNA binding assay protocol was adapted from previous work.^18^ Colocalization of ATTO 647N-labeled in-solution 60mer DNAs to Cy3B-labeled surface-bound 100mer DNAs tethered to the coverslip via a biotin-streptavidin linkage (Figure 1A,C). Flow cells were first coated with 0.2 mg/mL streptavidin (Millipore Sigma) in ELB. Excess streptavidin was washed away using ELB before adding 200 pM Cy3B-surface DNA to coat the surface and washing with ELB supplemented with oxygen scavenger (5 mM protocatechuic acid, 0.1 μM protocatechuate-3,4-dioxygenase) and redox pair (1 mM ascorbic acid, 1 mM methyl viologen). Colocalization was monitored in HSS extract supplemented with 5 nM unbound in-solution ATTO 647N-labeled DNA, 7.5 ng/μL nocodazole, 3 ng/μL linearized carrier DNA, 30 ng/μL carrier DNA, ATP regeneration system (ARS; 1.5 mM ATP, 7.5 mM Phosphocreatine, 0.01 mg/mL creatine phosphokinase), oxygen scavenger and redox pair. Reactions were monitored in a single field of view with continuous, alternating excitation between 532 nm and 642 nm with a 200 ms exposure time for 8 minutes. Experiments were repeated in two biological replicates.

### DNA circularization assay

The DNA circularization assay protocol was adapted from previous work.^18^ Flow cells were first coated with 0.2 mg/mL streptavidin (Millipore Sigma) in ELB. Excess streptavidin was washed away using ELB before adding 200 pM doubly labeled DNA substrate to coat the surface and washing with ELB supplemented with the oxygen scavenger and redox pair. End joining was then monitored in wild-type or depleted HSS extract supplemented with 3 ng/μL linearized carrier DNA, 30 ng/μL carrier DNA, 7.5 ng/μL nocodazole, ATP regeneration system, oxygen scavenger and redox pair. For add-back reactions, endogenous concentrations of proteins were added: 50 nM cdLigIV/XRCC4 and/or 50 nM XLF heterodimer. Reactions were monitored in a single field of view with stroboscopic, alternating excitation between 532 nm and 642 nm with a 200 ms exposure time imaged every 2.6 seconds for 15 minutes. Experiments were repeated in two biological replicates.

### LigIV DNA binding assay

Cy5-ybbR-LigIV/XRCC4 DNA binding was measured following a previous protocol.^17^ Flow cells were first coated with 0.2 mg/mL streptavidin (Millipore Sigma) and 10 mg/mL BSA (Sigma) in ELB. Excess streptavidin was washed away using ELB with 10 mg/mL BSA before adding 200 pM Cy3B-labeled surface DNA substrate to coat the surface and washing with ELB supplemented with 10 mg/mL BSA, the oxygen scavenger and redox pair. DNA binding was monitored in ELB with 20 nM Cy5-ybbR-LigIV/XRCC4, 1 mM ATP, oxygen scavenger and redox pair. Reactions were monitored over three fields of view with continuous, alternating excitation between 532 nm and 642 nm with a 200 ms exposure time for 3 minutes. Experiments were repeated in a biological replicate.

### Data analysis

Data were analyzed using a previously published automated pipeline in MATLAB.^54^ Briefly, cameras were aligned using a DNA standard containing Cy3B and ATTO 647N, movies were corrected for drift, bound DNA molecules were chosen based on their circularity, distance to nearest neighbor and intensity. Colocalization events were determined based on thresholds for ATTO 647N intensity and the distance between Cy3B and ATTO 647N foci, with thresholds set to minimize nonspecific binding at dark spots identified by the software (locations lacking surface-bound DNA). Positive colocalization events were required to last at least two frames and were allowed three frames between successive colocalization events to account for blinking. These events were quantified by either the time to first binding event, to determine the binding rate by fitting to a single exponential equation, or by their duration, to determine the dwell time of binding by fitting the dwell time histogram to a double-exponential decay. Confidence intervals for MLE exponential fits were determined by bootstrap resampling in MATLAB (MathWorks). Exact sample sizes are listed in their respective tables.

For FRET experiments, the total number of molecules was normalized to the total number of colocalized molecules in each field of view. FRET data were analyzed using the same software package as the colocalization data. For visualization, background-corrected fluorescence intensities were analyzed in MATLAB (MathWorks) to correct for bleed-through from the green channel to the red channel, correct for ATTO 647N excitation from the green laser, gamma correct for Cy3B-ATTO 647N and truncated after photobleaching. Events showing FRET were fit in the same manner as colocalization events.

## Acknowledgements

We thank current and former members of the Loparo laboratory and Johannes Walter for discussions related to the manuscript, Andrew Moreno and Jinghang Zhang for help in learning the long-range complex assay.

## Author contributions

Brandon C. Case: Designed the project and experiments, prepared *Xenopus* egg extract, performed antibody purifications, gel-based assays, LRC and SRC single-molecule experiments, data analysis, and writing – original draft, review, and editing. Isabel T. Burger: Performed LRC and SRC single-molecule experiments and data analysis. Joseph J. Loparo: Designed the project and experiments, writing – review and editing.

## Data availability

Code for the initial single-molecule imaging pipeline was previously published.^54^ Single-molecule traces, uncropped images, replicates of gel data, and downstream analysis code from this manuscript are available on Zenodo at DOI: 10.5281/zenodo.21907804

