## Supplemental Figure 1 for "Core NHEJ Factors Maintain Synapsis Across Diverse DNA End Structures"

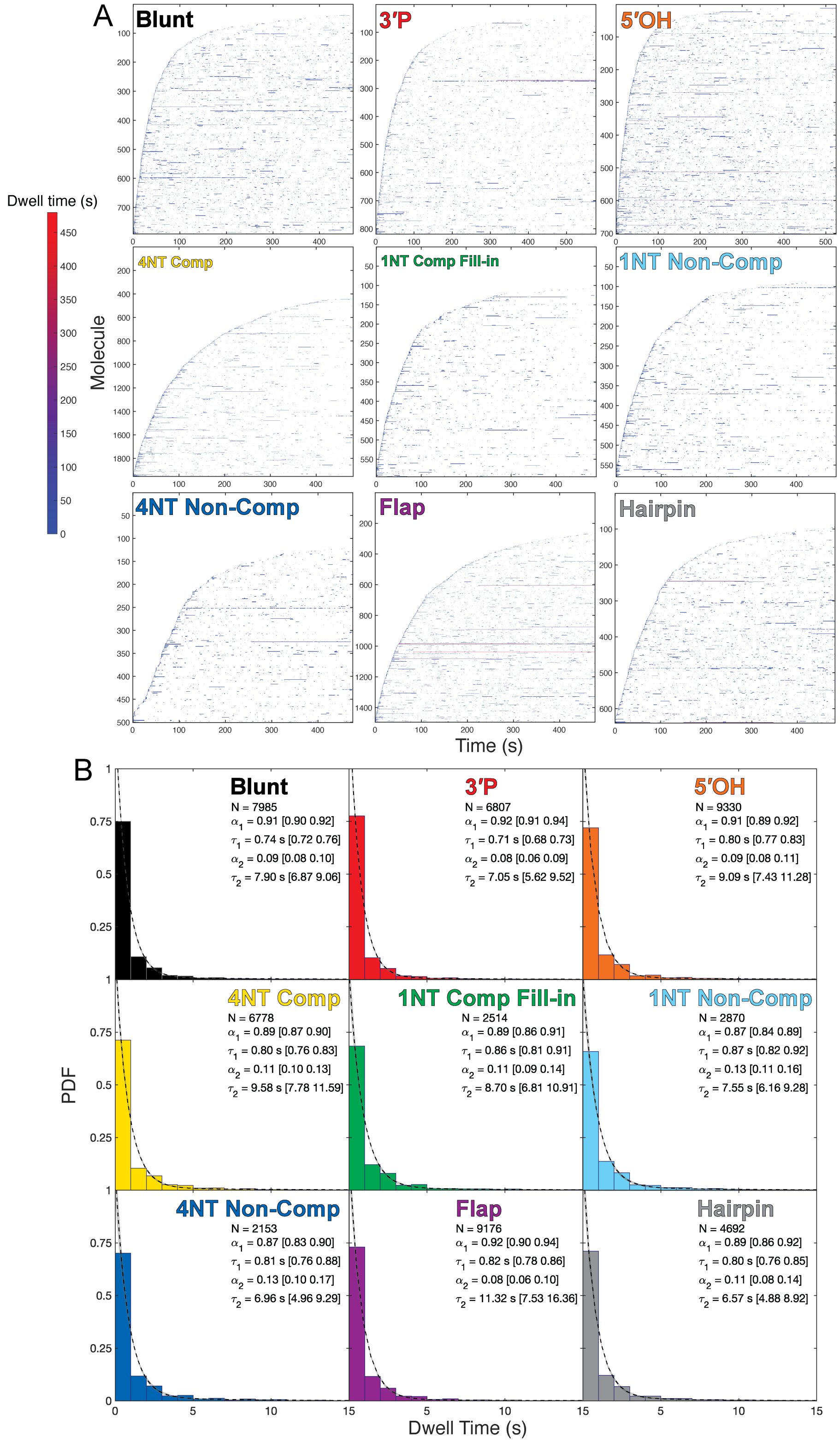

**Supplemental Figure 1**: (A) Rastergrams from the two-piece single molecule colocalization assay testing LRC formation and stability using *Xenopus* HSS extract for each DNA substrate. Individual molecules were sorted by time to first binding and shaded based on the dwell time of each DNA binding event. (B) Dwell time histograms representing the probability density function (PDF) for each DNA type; fits shown in a dashed black line with confidence intervals shaded in grey.

**Supplemental Figure 2**:

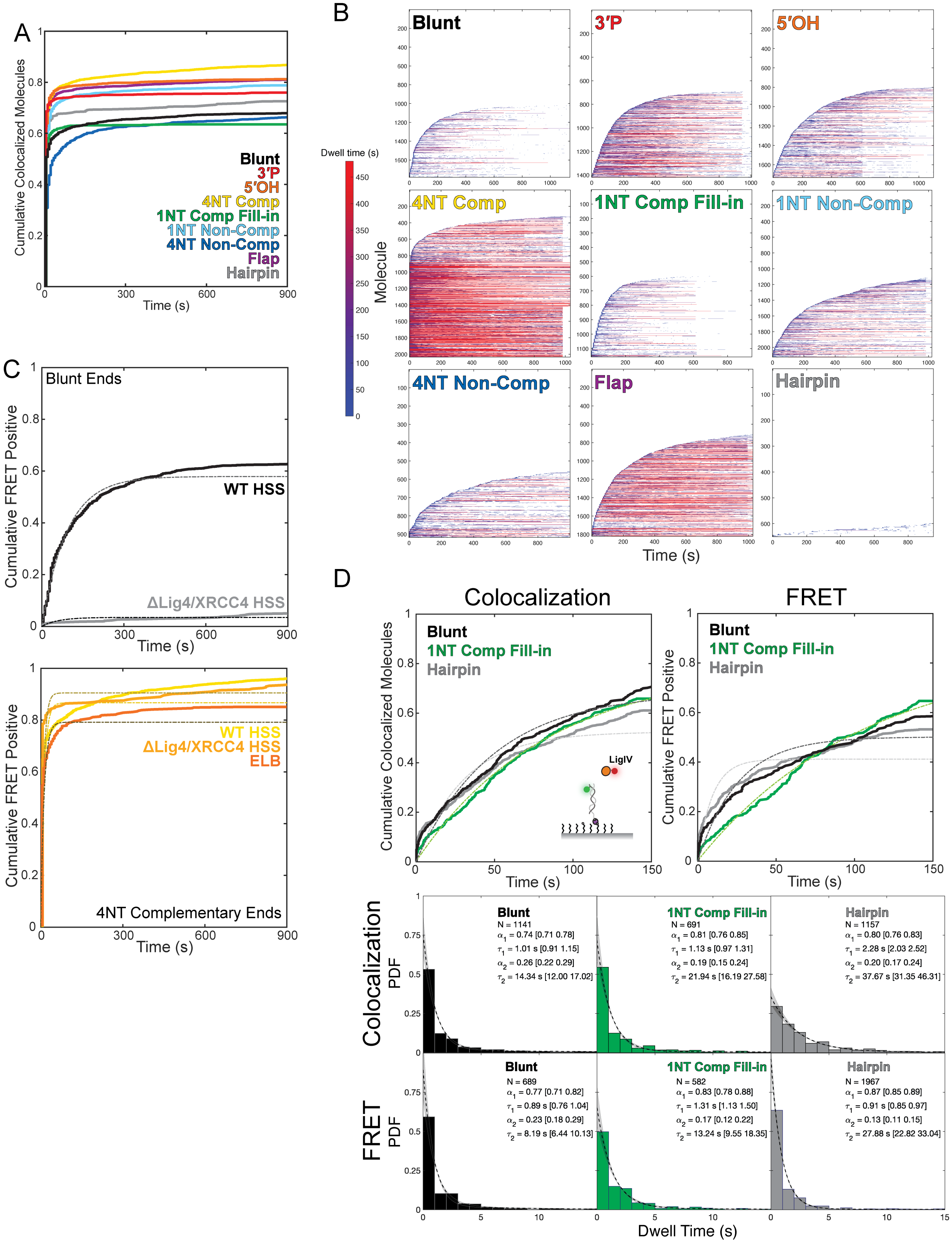

**Supplemental Figure 2**: (A) Representative plot comparing cumulative percent colocalized molecules for each DNA type corresponding to the total number of DNA molecules able to undergo FRET for each experiment. (B) Rastergrams from the smFRET circularization assay testing SRC formation using wild type *Xenopus* HSS extract. Individual molecules were sorted by time to first FRET and shaded based on the dwell time of each DNA binding event. (C) Cumulative percent FRET positive curves testing SRC formation in different conditions for blunt (upper) and 4NT complementary (lower) ends. (D) Comparison of Cy5-ybbR-LigIV/XRCC4 binding to various DNA types by both colocalization and FRET with dwell time histograms shown to compare the stability of binding interactions.

**Supplemental Figure 3**:

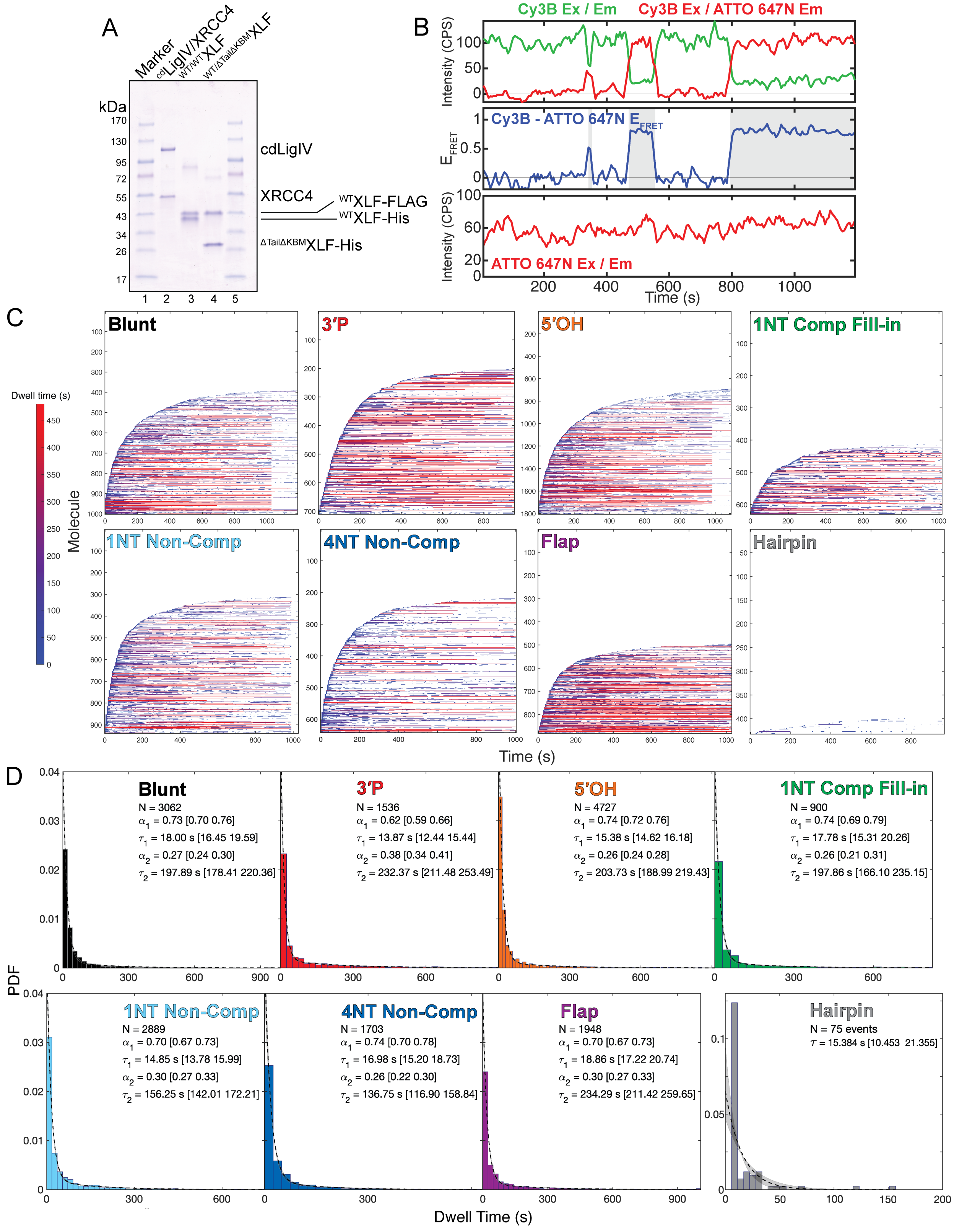

**Supplemental Figure 3**: (A) Protein purification gel showing recombinant proteins used in this study. (B) Representative single-molecule trace showing DNA stably bound on the surface with both Cy3B (upper) and ATTO 647N (lower) over time. A decrease in Cy3B emission and concomitant increase in ATTO 647N emission upon Cy3B excitation (upper) are indicative of FRET (shaded area, middle). (C) Rastergrams from the smFRET circularization assay testing SRC formation and stability using ΔXRCC4/LigIV *Xenopus* HSS extract with add-back of $^{cd}$LigIV/XRCC4 for different DNA substrates. Individual molecules were sorted by time to first FRET and shaded based on the dwell time of each DNA binding event.  (D) Dwell time histograms representing the probability density function (PDF) for each DNA substrate; fits shown in a dashed black line with confidence intervals shaded in grey.

**Supplemental Figure 4**:

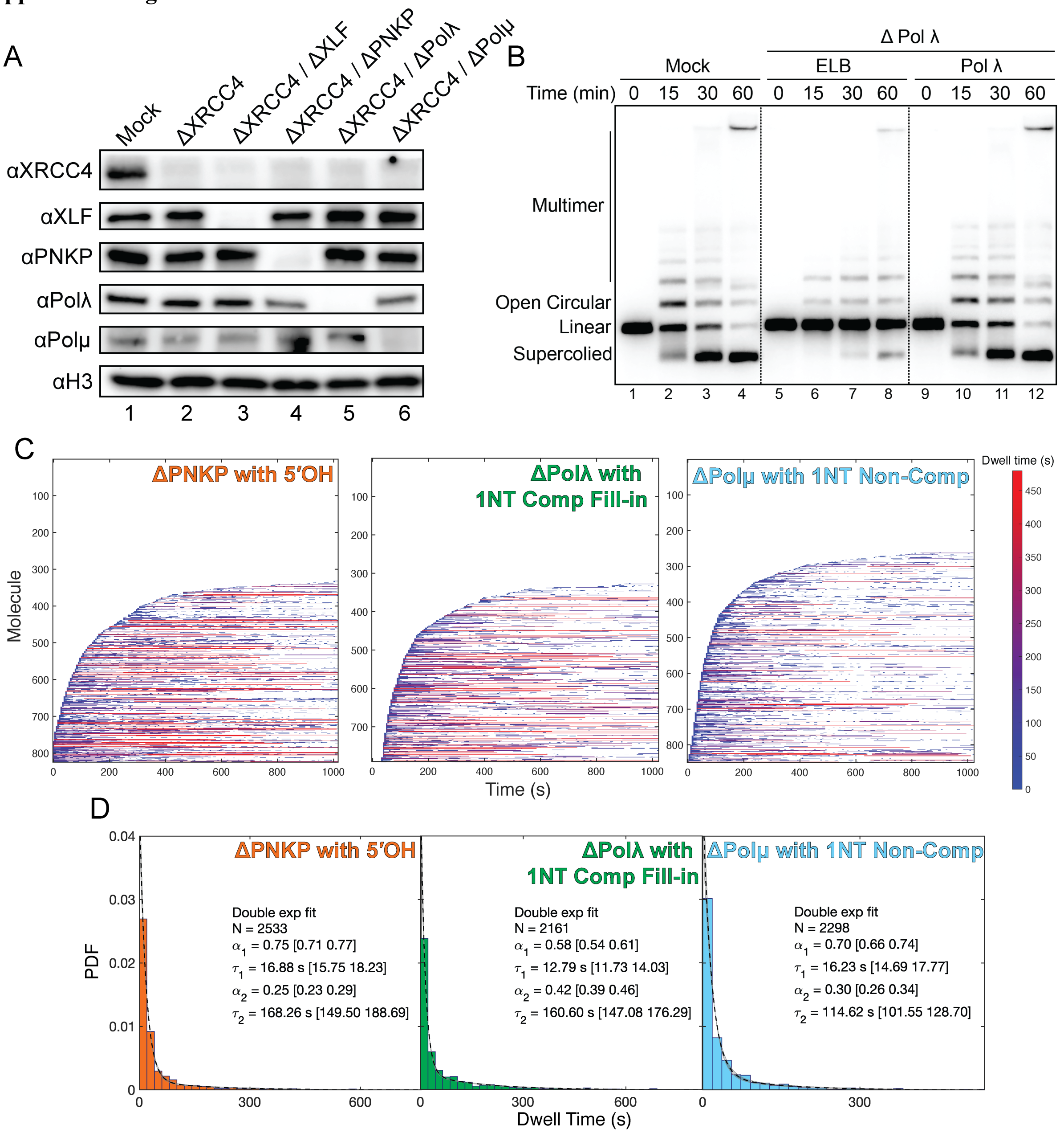


**Supplemental Figure 4**: (A) Western blots showing successful depletion of different proteins used in this work. (B) Bulk end-joining assay showing the depletion and add-back of Pol λ on a 1NT Comp Fill-in substrate. (C) Rastergrams from the smFRET circularization assay testing SRC formation and stability using ΔXRCC4/LigIV *Xenopus* HSS extract with add-back of $^{cd}$LigIV/XRCC4 and additional depletion of various processing factors on their cognate DNA substrates. Individual molecules were sorted by time to first FRET and shaded based on the dwell time of each DNA binding event. (D) Dwell time histograms representing the probability density function (PDF) for each condition; fits shown in a dashed black line with confidence intervals shaded in grey.

**Supplemental Figure 5**:

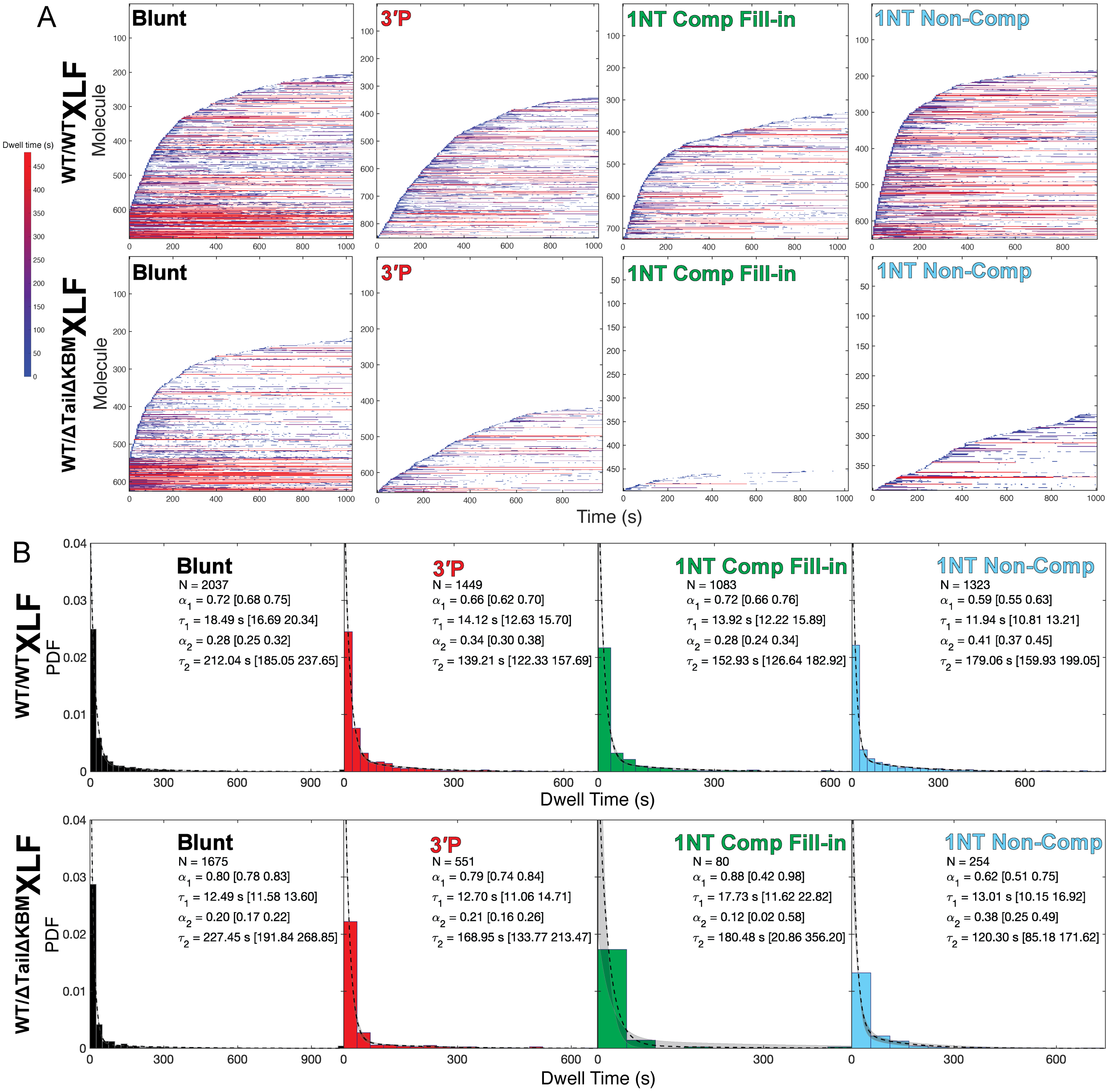


**Supplemental Figure 5**: (A) Rastergrams from the smFRET circularization assay testing SRC formation and stability using ΔXLF/ΔXRCC4/LigIV *Xenopus* HSS extract with add-back of $^{cd}$LigIV/XRCC4 and XLF heterodimers for various DNA substrates. Individual molecules were sorted by time to first FRET and shaded based on the dwell time of each DNA binding event. (B) Dwell time histograms representing the probability density function (PDF) for each condition; fits shown in a dashed black line with confidence intervals shaded in grey.

**Supplementary Table 1** – Oligos for DNA End Structures

| DNA End | Oligo | Label | PNK Rxn | 5' to 3' sequence | Pairing for SphI annealing | Pairing for AatII annealing | DNA End |
|---|---|---|---|---|---|---|---|
| Blunt | oBC54 | $^{32}P$ | Yes | CGT ACC GCT CTA TGC | oBC54/58 - 5' CGT ACC GCT CTA TGC<br>oBC55 - 3' G TAC GCA TGG CGA GAT ACG | TCC GAG AAT ACC GCC ACG T 3' - oBC56<br>AGG CTC TTA TGG CGG 5' - oBC57/59 | Blunt |
| | oBC55 | - | Yes | GCA TAG AGC GGT ACG CAT G | | | |
| | oBC56 | - | Yes | TCC GAG AAT ACC GCC ACG T | | | |
| | oBC57 | $^{32}P$ | Yes | GGC GGT ATT CTC GGA | | | |
| | oBC58 | Cy3B | Yes | CGT ACC GCT C/iAmMC6T/A TGC | | | |
| | oBC59 | ATTO 647N | Yes | GGC GGT ATT C/iAmMC6T/C GGA | | | |
| 3'P | oBC55 | - | Yes | GCA TAG AGC GGT ACG CAT G | oBC62 - 5' CGT ACC GCT CTA TGC<br>oBC55 - 3' G TAC GCA TGG CGA GAT ACG | TCC GAG AAT ACC GCC ACG T 3' - oBC56<br>AGG CTC TTA TGG CGG 5' - oBC63 | 3'P |
| | oBC56 | - | Yes | TCC GAG AAT ACC GCC ACG T | | | |
| | oBC62 | Cy3B | No | /5Phos/CGT ACC GCT C/iAmMC6T/A TGC/3Phos/ | | | |
| | oBC63 | ATTO 647N | No | /5Phos/GGC GGT ATT C/iAmMC6T/C GGA/3Phos/ | | | |
| 5'OH | oBC55 | - | No | GCA TAG AGC GGT ACG CAT G | oBC58 - 5' CGT ACC GCT CTA TGC<br>oBC55 - 3' G TAC GCA TGG CGA GAT ACG | TCC GAG AAT ACC GCC ACG T 3' - oBC56<br>AGG CTC TTA TGG CGG 5' - oBC59 | 5'OH |
| | oBC56 | - | No | TCC GAG AAT ACC GCC ACG T | | | |
| | oBC58 | Cy3B | Yes | CGT ACC GCT C/iAmMC6T/A TGC | | | |
| | oBC59 | ATTO 647N | Yes | GGC GGT ATT C/iAmMC6T/C GGA | | | |
| 4NT Comp | oBC58 | Cy3B | Yes | CGT ACC GCT C/iAmMC6T/A TGC | oBC58 - 5' CGT ACC GCT CTA TGC<br>oBC97 - 3' G TAC GCA TGG CGA GAT ACG CTAG | GATC TCC GAG AAT ACC GCC ACG T 3' - oBC98<br>AGG CTC TTA TGG CGG 5' - oBC59 | 4NT Comp |
| | oBC59 | ATTO 647N | Yes | GGC GGT ATT C/iAmMC6T/C GGA | | | |
| | oBC97 | - | Yes | GAT CGC ATA GAG CGG TAC GCA TG | | | |
| | oBC98 | - | Yes | GAT CTC CGA GAA TAC CGC CAC GT | | | |
| 1NT Comp Fill-in | oBC54 | $^{32}P$ | Yes | CGT ACC GCT CTA TGC | oBC54/58 - 5' CG TAC CGC TCT ATG C<br>oBC107 - 3' GTA CGC ATG GCG AGA TAC GAC | GTC GTA ACG CCC TAT GCA CGT 3' - oBC108<br>G CAT TGC GGG ATA CG 5' - oBC66/67 | 1NT Comp Fill-in |
| | oBC58 | Cy3B | Yes | CGT ACC GCT C/iAmMC6T/A TGC | | | |
| | oBC66 | $^{32}P$ | Yes | GCA TAG GGC GTT ACG | | | |
| | oBC67 | ATTO 647N | Yes | GCA TAG GGC G/iAmMC6T/T ACG | | | |
| | oBC107 | - | Yes | CAG CAT AGA GCG GTA CGC ATG | | | |
| | oBC108 | - | Yes | GTC GTA ACG CCC TAT GCA CGT | | | |
| 1NT Non-Comp | oBC58 | Cy3B | Yes | CGT ACC GCT C/iAmMC6T/A TGC | oBC58 - 5' CG TAC CGC TCT ATG C<br>oBC276 - 3' GTA CGC ATG GCG AGA TAC GG | GC GTA ACG CCC TAT GCA CGT 3' - oBC275<br>G CAT TGC GGG ATA CG 5' - oBC67 | 1NT NonComp |
| | oBC67 | ATTO 647N | Yes | GCA TAG GGC G/iAmMC6T/T ACG | | | |
| | oBC275 | - | Yes | GCG TAA CGC CCT ATG CAC GT | | | |
| | oBC276 | - | Yes | GGC ATA GAG CGG TAC GCA TG | | | |
| 4NT Non-Comp | oBC58 | Cy3B | Yes | CGT ACC GCT C/iAmMC6T/A TGC | oBC58 - 5' CG TAC CGC TCT ATG C<br>oBC287 - 3' GTA CGC ATG GCG AGA TAC GGGGG | GGGGC GTA ACG CCC TAT GCA CGT 3' - oBC288<br>G CAT TGC GGG ATA CG 5' - oBC67 | 4NT NonComp |
| | oBC67 | ATTO 647N | Yes | GCA TAG GGC G/iAmMC6T/T ACG | | | |
| | oBC287 | - | Yes | GGG GGC ATA GAG CGG TAC GCA TG | | | |
| | oBC288 | - | Yes | GGG GCG TAA CGC CCT ATG CAC GT | | | |
| Flap | oBC58 | Cy3B | Yes | CGT ACC GCT C/iAmMC6T/A TGC | oBC58 - 5' CGT ACC GCT CTA TGC<br>oBC70 - 3' G TAC GCA TGG CGA GAT ACG GGC CAA AA | AAA ACC GGT CCG AGA ATA CCG CCA CGT 3' - oBC71<br>A GGC TCT TAT GGC GG 5' - oBC59 | Flap |
| | oBC59 | ATTO 647N | Yes | GGC GGT ATT C/iAmMC6T/C GGA | | | |
| | oBC70 | - | Yes | AAA ACC GGG CAT AGA GCG GTA CGC ATG | | | |
| | oBC71 | - | Yes | AAA ACC GGT CCG AGA ATA CCG CCA CGT | | | |
| Hairpin | oBC270 | Cy3B | Yes | GAT CTG GCC TG/iAmMC6T/ CTT ATA AGA CAG GCC AGA TCC ATG | oBC270 - 5' GAT CTG GCC TGT CTT [illegible]<br>G TAC CTA GAC CGG ACA GA [illegible] | [illegible] GGA CGC AAG ACA TCT ACG T<br>[illegible] CT GCG TTC TGT AGA 5' - oBC271 | Hairpin |
| | oBC271 | ATTO 647N | Yes | AGA TGT CTT GCG /iAmMC6T/CC TAG GAC GCA AGA CAT CTA CGT | | | |

Note: Nucleotides in red are flourescently labeled through a /iAmMC6T/ linkage from IDT; nucleotides in green are radiolabeled with $^{32}P$; nucleotides in purple are phosphorlated from IDT through /5Phos/ or /3Phos/ modification; nucleotides in orange are non-phosphorylated from exclusion in the PNK reaction; nucleotides in blue are complementary between strands.

**Supplementary Table 2** – Oligos for DNA backbones and standards

| Purpose | Oligo | Label | 5' to 3' sequence |
|---|---|---|---|
| DNA Standard - Biotin | oBC129 | - | /5Biosg/ACT CAA CCA AGT CAT TCT GAG AAT AGT GTA TGC GGC G |
| DNA Standard - Cy3B | oBC130 | Cy3B | /5AmMC6/CCG AGT TGC TCT TGC CCG GCG TCA A |
| DNA Standard - ATTO 647N | oBC131 | ATTO 647N | CGG GAT AAT ACC GCG CCA CAT AGC AGA AC/3AmMO/ |
| DNA Standard Backbone | oBC134 | - | AGA TCC TTG AGA GTT TTC GCC CCG AAG AAC GTT TTC CAA TGA TGA GCA CTT TTA AAG TTC TGC TAT GTG GCG CGG TAT TAT CCC GTA TTG ACG CCG GGCAAG AGC AAC TCG GTC GCC GCA TAC ACT ATT CTC AGA ATG ACT TGG TTG AGT |
| LRC Bound DNA Backbone | oIB06 | - | GAG ACG TCT GCC TGT TGA ACA AGG TAC CCA GCT TTT GTT CCC TTT AGT GAG GGT TAA TTG/3Bio/ |
| LRC Bound DNA Backbone | oIB07 | - | CAA TTA ACC CTC ACT AAA GGG AAC AAA AGC TGG GTA CCT TGT TCA ACA GGC AGA CGT CTC ACG T |
| LRC In-solution DNA Backbone | oIB08 | - | GGG CGA CTC ACT ATA GGG CGA ATT GGA GCT CAT AAT GTC GGG CAA TCA GGT GCG ACA ATC TAT CGA TTG TAT GGG AAG CCC GAT GCG CCA GGC TTC CAT G |
| LRC In-solution DNA Backbone | oIB09 | - | GAA GCC TGG CGC ATC GGG CTT CCC ATA CAA TCG ATA GAT TGT CGC ACC TGA TTG CCC GAC ATT ATG AGC TCC AAT TCG CCC TAT AGT GAG TCG CCC/3Bio/ |
| SRC Biotinylation Strand | oBC127 | - | TCA GCC TGC AGG AA/iBiodT/ TCG ATA TCC C |
